# Resurgent Na^+^ currents promote ultrafast spiking in projection neurons that drive fine motor control

**DOI:** 10.1101/2021.09.09.459677

**Authors:** Benjamin M. Zemel, Alexander A. Nevue, Andre Dagostin, Peter V. Lovell, Claudio V. Mello, Henrique von Gersdorff

## Abstract

The underlying mechanisms that promote precise spiking in upper motor neurons controlling fine motor skills are not well understood. Here we report that projection neurons in the adult zebra finch song nucleus RA display: 1) robust high-frequency firing, 2) ultra-short half-width spike waveforms, 3) superfast Na^+^ current inactivation kinetics and 4) large resurgent Na^+^ currents (I_NaR_). These spiking properties closely resemble those of specialized pyramidal neurons in mammalian motor cortex and are well suited for precise temporal coding. They emerge during the critical period for vocal learning in males but not females, coinciding with a complete switch of modulatory Na^+^ channel subunit expression from Navβ3 to Navβ4. Dynamic clamping and dialysis of Navβ4’s C-terminal peptide into juvenile RA neurons provide evidence that this subunit, and its associated I_NaR_, promote neuronal excitability. We propose that Navβ4 underpins I_NaR_ that facilitates precise, prolonged, and reliable high-frequency firing in upper motor neurons.

## Introduction

The speed and accuracy of muscle control and coordination depends on the spiking activity of upper motor neurons^1^. In mammals, a subclass of layer 5 pyramidal neurons (L5PNs) project from the motor cortex to various targets in the brainstem and spinal cord. These cells are involved in specific aspects of fine motor control and they produce narrow half-width action potentials (APs)^2,3^. Changes to their intrinsic properties have been implicated in facilitating the learning of complex motor skills in some species^4-8^. Notably, primates and cats possess varying numbers of very large L5PNs with wide-caliber myelinated axons, fast AP conduction velocities and ultra-narrow AP spikes^9-11^. These specialized Betz-type cells, first discovered in primates, send projections that often terminate directly onto lower motor neurons^7^, and are thought to be involved in highly refined aspects of motor control^1,5,9,10^.

Birds display a diverse array of complex behaviors and cognitive skills ranging from elaborate nest building and tool usage to episodic memory and vocal mimicry^12-14^. Remarkably, birds accomplish this without a typical six-layered neocortex, which underpins the capacity for complex motor skills in mammals^15,16^. Nonetheless, avian pallial nuclei form microcircuits that appear analogous to those in the mammalian neocortex^15-19^. In songbirds, the robust nucleus of the arcopallium (RA) plays key roles in singing and provides direct descending projections to brainstem motor neurons that innervate the avian vocal organ (syrinx; Fig. 1a)^20-22^ and respiratory muscles^23^. RA projection neurons (RAPNs) can thus be considered analogous to L5PNs in motor cortex^16-19^. Indeed, these cells share important features, like wide-caliber myelinated axons and multiple spine-studded basal dendrites^24^, although RAPNs lack the large, multilayer-spanning apical tufted dendrites that are a hallmark of L5PNs^25-27^. However, detailed knowledge of the ion channel composition, biophysical properties and firing patterns of RAPNs is limited. Therefore, it is still unclear to what extent they function in an analogous manner to L5PNs.

**Figure 1.**
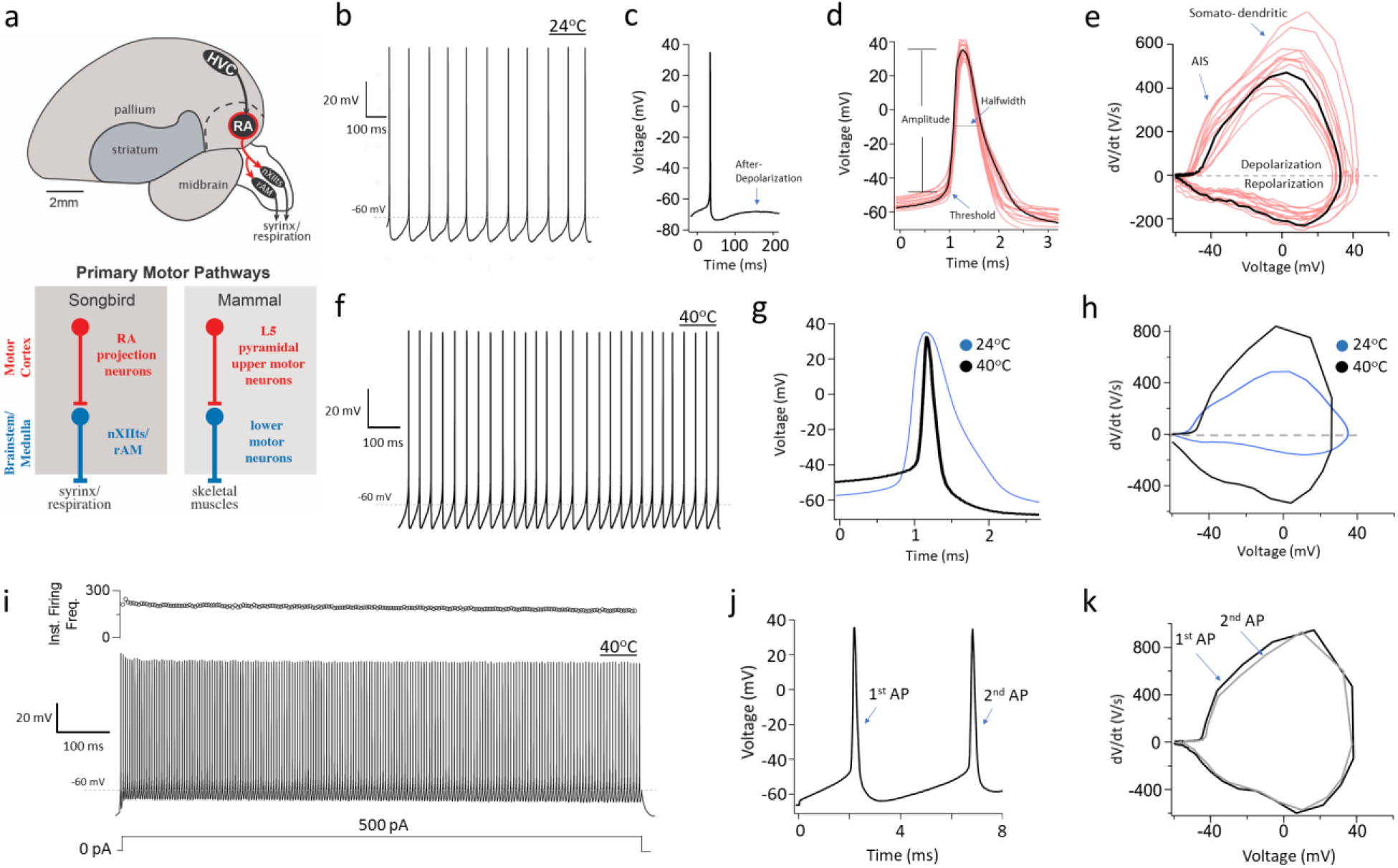
Intrinsic excitable properties of RA projection neurons (RAPNs) in adult male zebra finches at room and physiological temperatures. **a**. Top: Zebra finch vocal control circuitry; nuclei and connections of the direct vocal-motor pathway shown in black, RA and its descending projections highlighted in red, other nuclei and connections omitted for clarity. Dashed line represents the dorsal boundary of the arcopallium. HVC is a proper name; RA, robust nucleus of the arcopallium; nXIIts, tracheosyringeal division of the hypoglossal nerve nucleus; RAm, nucleus retroambigualis medialis. Bottom: Diagram depicting upper (red) and lower (blue) motor neurons of the descending motor pathway in zebra finch, a songbird (left), and mammals (right). **b**. Whole cell current clamp recording of spontaneous APs during 1 sec. **c**. Spontaneous AP from a different cell than (b), arrow indicates after-depolarization. **d**. Representative spontaneous AP from (b; black) with overlay of spontaneous APs from an additional 15 RAPNs (pink); values for the indicated parameters are presented in Table 1. **e**. Phase plane plots of the AP in (d; black) and from another 15 RAPNs (pink, same cells as in d); indicated are the presumed contributions of the axonal initial segment (AIS) and somato-dendritic components to the biphasic depolarization component in the phase plane plot. **f**. Whole cell current clamp recording of spontaneous APs during 1 sec in a RAPN at physiological temperature (40°C). **g**. Representative spontaneous AP recorded at physiological temperature (40°C; black) with overlay of a spontaneous AP recorded at room temperature (24°C; blue). **h**. Overlayed phase plane plots of the spontaneous APs from g. **i**. Representative AP train elicited by a 1 sec 500 pA current injection at 40°C; the corresponding plot of instantaneous firing frequency as function of time is shown at the top. **j**. First two APs from (i). **k**. Overlay of the phase plane plots from the two APs in (j).

Zebra finch RAPNs face considerable spiking demands during singing, which requires superfast, temporally precise coordination of syringeal and respiratory musculature^28-30^. As the adult male sings, RAPNs exhibit remarkably precise spike timing (variance ∼0.23 ms)^31^. RAPNs also exhibit increased burstiness during the developmental song learning period, their instantaneous firing rates changing from 100-200 Hz when they produce immature vocalizations (subsong) to 300-600 Hz when song becomes mature (crystallized). Average overall spike rates of RAPNs increase from 36 Hz at subsong to 71 Hz in adults^32^. Song maturation thus correlates with reduced variability in the timing of increasingly high frequency bursts with a refinement of single spike firing precision. Importantly, nerve firing rates of >75 Hz are required for force summation in the superfast syrinx muscles^33^. This high spike frequency in RAPNs during song production is energetically demanding as indicated by increased staining for cytochrome C in maturing males, but not female zebra finches^34^.

Songbird RAPNs thus seem to share with Betz-type L5PNs similar evolutionary pressure for fast and precise signaling, a constraint that can lead to neurons in unrelated species sharing similar expression patterns for a specific repertoire of ion channels. A good example are electric fish species from different continents, which have independently evolved electrocytes exhibiting similar physiology and molecular convergence^35,36^. Indeed, high-frequency firing electrocytes with extremely narrow AP spikes co-express fast activating Na^+^ and K^+^ channels^37,38^.

Here we explored the excitability of zebra finch RAPNs, posing the following question: Are the properties of Na^+^ currents in RAPNs specialized to produce narrow, non-adapting spikes that enable precise high-frequency burst firing? We demonstrate that RAPNs share strikingly similar spiking properties and ion channel subunits with Betz-type L5PNs. We provide strong evidence linking the presence of voltage-gated Na^+^ channel auxiliary β4 subunit (Navβ4) to large resurgent Na^+^ currents (I_NaR_) in RAPNs. We also provide compelling evidence that the ability of maturing RAPNs to produce narrow APs strongly correlates with increasing Navβ4 mRNA expression during the critical period for vocal learning in males, but not females that do not sing. We conclude that Navβ4 and related I_NaR_ likely play an important role in fine tuning the intrinsic excitability of RAPNs. We propose these biophysical properties allow a subclass of large upper motor neurons to operate as highly specialized control units for the production of rapid and precise movements.

## Results

### Spontaneous and evoked spiking in RAPNs

Whole-cell current clamp recordings performed in adult male zebra finch brain slices revealed that RAPNs fire spontaneously, with tonic activity in the absence of synaptic input. Action potential (APs) spikes had large amplitudes and narrow, submillisecond durations with a prominent after-depolarization that facilitated the production of subsequent spikes (Fig. 1b-d; spikes at ∼24°C; see Table 1). Recorded RAPNs were easily distinguished from local inhibitory GABAergic interneurons based on differences in their spiking properties and morphology (Supplementary Fig. 1). These results are consistent with previous studies^24,39-42^. However, phase plane plots from the first derivative of these spontaneous APs revealed a biphasic AP upstroke, (Fig. 1e), which has not been previously described in songbirds. This biphasic AP is a hallmark of L5PNs in mammalian neocortex and suggests a high density of voltage-gated Na^+^ channels (Nav) at the axon initial segment^43^. The phase plots also revealed large maximum rates of depolarization and repolarization (Fig. 1e; Table 1), which likely facilitates high frequency firing during *in vivo* song production^32^.

At physiological avian body temperature (∼40°C^44^) the spontaneous firing frequency increased four-fold compared to room temperature (Fig. 1b and f; Table 1;^45^). Importantly, the AP waveform had a smaller amplitude at 40°C and an extremely narrow half-width (∼0.2 ms; Fig. 1g; Table 1). The APs also exhibited faster maximum rates of depolarization and repolarization in the phase plot at 40°C (Fig. 1h; Table 1). During positive current injections adult RAPNs produced high-frequency AP trains at 40°C (Fig. 1f). Large amplitude spikes were triggered for the duration of the current injection with only a modest initial adaptation of spike amplitude. In fact, RAPNs showed little change in the maximum rate of depolarization between the 1^st^ and 2^nd^ spikes, despite a faster instantaneous firing frequency (IFF) at 40°C (Fig. 1g; for ∼24°C data see Supplementary Fig. 2). We also note that during negative current injections at room temperature, we observed a voltage sag that is consistent with HCN-mediated currents (5.0 ± 0.68 mV between the peak and steady-state voltage during a −150 pA step stimulus), as previously described for RAPNs^40^.

**Figure 2.**
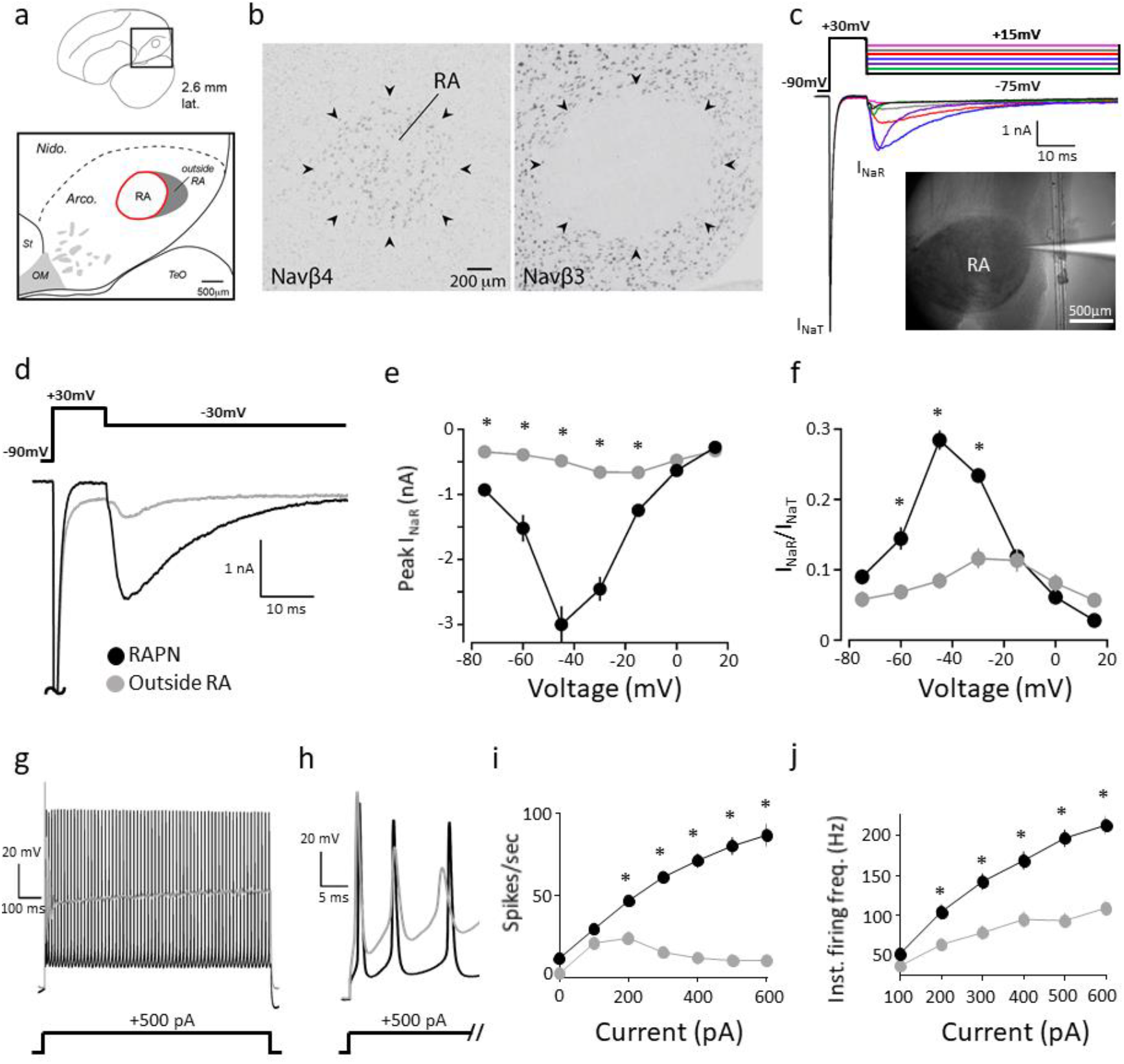
Expression of Navβ4 and β3 mRNAs, I_NaR_ and intrinsic excitability in the arcopallium of adult male zebra finches. **a**. Anatomical drawing indicating the position of RA within the arcopallium in a parasagittal section; distance from the midline indicated; the shaded area represents the region outside RA that is compared with recordings within RA. Arco., arcopallium; Nido., Nidopallium; OM, occipitomesencephalic tract; St, striatum; TeO, Optic tectum. orientation: dorsal is up and anterior to the left. **b**. *In situ* hybridization images show high Navβ4 expression (left) and lack of Navβ3 expression (right) in RA (indicated by arrowheads). **c**. Representative examples of I_NaT_ and I_NaR_ elicited in an RAPN by the voltage clamp protocol at the top. I_NaT_ was elicited via a 10 ms step to +30 mV followed by test potentials to +15 (pink), 0 (grey), −15 (red), −30 (blue), −45 (purple), −60 (green) and −75 mV (black) to elicit I_NaR_. The cell was held at −90 mV during a 2 sec intersweep interval. Inset: Image of a patch pipette filled with fluorescent dye for RA recordings. **d**. Detail views of example I_NaT_ and I_NaR_ elicited in an RAPN (black) and in a neuron outside RA (grey) by the voltage clamp protocol on the top. The I_NaT_ peaks have been truncated. **e**. Average I-V curves for the peak I_NaR_ in RAPNs (black) and in neurons outside RA (grey) (two-way ANOVA with Tukey’s post hoc; P < 0.0001, F (6, 146) = 27.00, N = 12 RAPNs and 11 neurons outside RA). **f**. Average I-V curves after normalization of the peak I_NaR_ to the peak I_NaT_ measured in a given sweep then averaged across cells (two-way ANOVA with Tukey’s post hoc; P < 0.0001, F (6, 147) = 29.36, N = 12 RAPNs and 11 neurons outside RA). **g**. Overlay of AP trains elicited by a 1 sec 500 pA current injection in an RAPN and in a neuron outside RA. **h**. The first 3 APs from the AP trains shown in (g). **i**. Average elicited spikes/sec as a function of current injected (two-way ANOVA with Tukey’s post-hoc; P < 0.0001, F (6, 160) = 22.79, N = 18 RAPNs and 12 neurons outside RA). **j**. Average instantaneous firing frequencies measured as a function of current injected (two-way ANOVA with Tukey’s post-hoc; P < 0.0001, F (5, 138) = 5.647, N = 18 RAPNs and 12 neurons outside RA). For data in e-f and i-j, * = P ≤ 0.05 by post-hoc analyses.

These results show that RAPNs have similar electrophysiological properies to cortical L5PNs that also display spontaneous APs with sub-millisecond half-widths, temperature-sensitive amplitudes, and modestly adapting AP firing during positive current injections^43,46-48^. Importantly, the prolonged high frequency spiking of RAPNs suggests a high degree of availability of Nav channels. We therefore hypothesized that adult RAPNs contain a large pool of Nav channels and specialized mechanisms that limit their dwell time in non-conducting inactivated states.

### Navβ4 expression predicts a robust resurgent Na^+^ current (I_NaR_)

Recently, we have identified several transcripts for voltage-gated Na^+^ channel beta (Navβ) auxiliary subunits as either positive (Navβ4) or negative (Navβ3) markers of adult male RA in the arcopallium^49^. While the functional role of these different Navβ subunits in RA neuronal excitability is unknown, Navβ4 is of particular interest because it promotes narrow spike waveforms and high frequency firing via a resurgent sodium current (I_NaR_)^50-52^. We therefore hypothesized that Navβ4 might be a key determinant of excitability properties in RA. High expression of Navβ4 mRNA is restricted to RA in the arcopallium of adult male zebra finches^49^, contrasting sharply with Navβ3, which is all but absent (Fig. 2a-b). Navβ4 is thought to promote high frequency firing by limiting classical inactivation of Nav channels and promoting I_NaR_^53^. Based on its expression levels, we predicted that neurons within RA would have a larger I_NaR_ than those in the arcopallium outside RA. We thus performed whole-cell voltage clamp recordings in RA sagittal slices from adult male finches (Fig. 2c). Voltage steps from −90 mV to +30 mV elicited large transient sodium currents (I_NaT_) and, as predicted, subsequent steps to a range of test potentials (+15 mV to −75 mV) yielded robust I_NaR_ (Fig. 2c) with a peak of −3.0 nA ± 0.28 (Mean ± SEM) at the −45 mV test potential (Fig. 2e). The decay kinetics of I_NaR_ was voltage dependent, with single exponential decay time constants ranging from 2 to 45 ms (Supplementary Fig. 3a-b). These results reveal an exceptionally large I_NaR_ in RAPNs compared to those recorded in mammalian^52^ and chicken brainstem neurons^54^.

**Figure 3.**
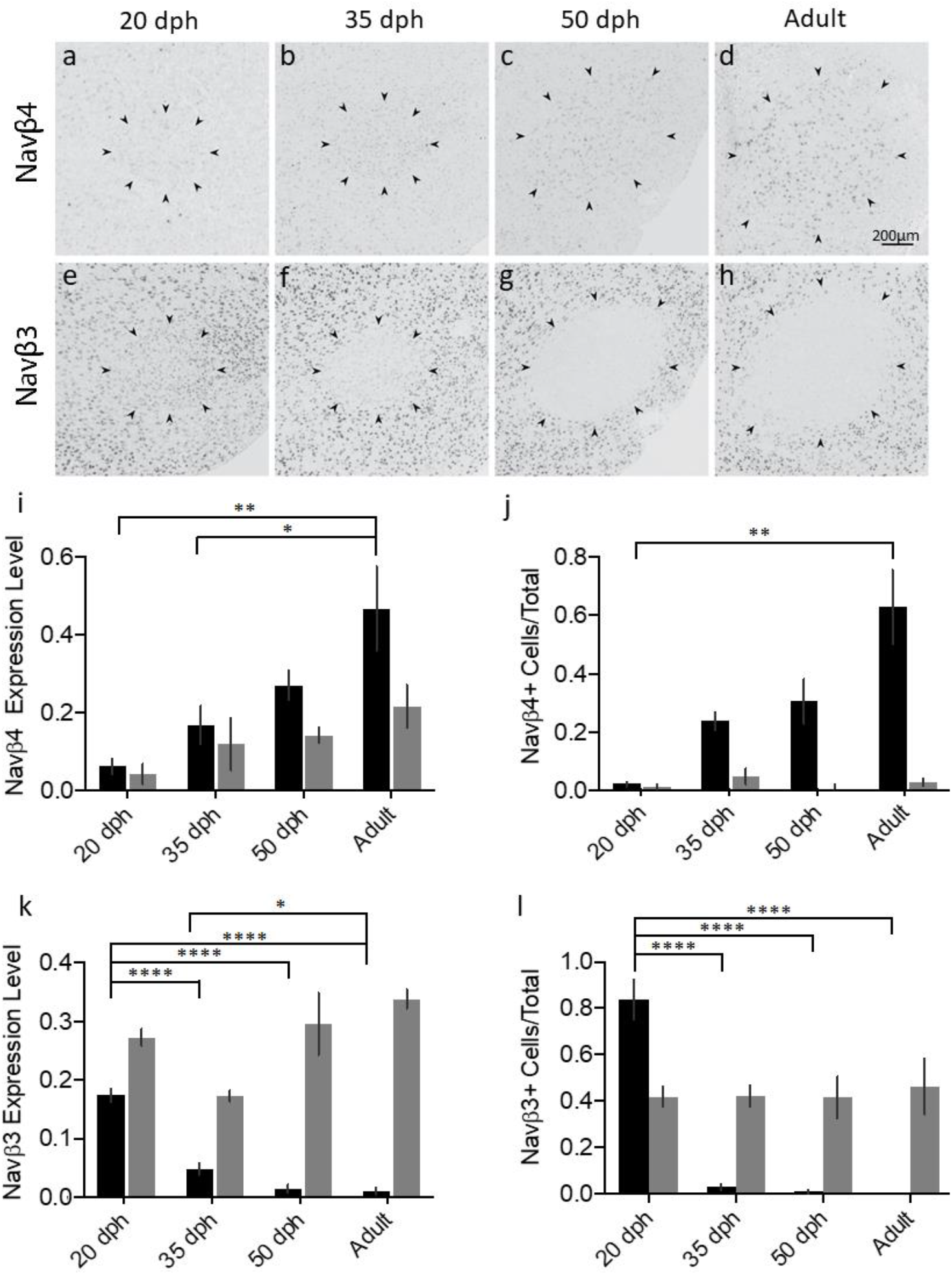
Age-dependent changes in expression of Navβ3 and 4 mRNAs in the arcopallium of male zebra finches across ages. **a-d**. Representative *in situ* hybridization images for Navβ4 mRNA within RA across ages; black arrowheads indicate RA borders. **e-h**. Representative *in situ* hybridization images for Navβ3 mRNA within RA across ages. Black arrowheads indicate RA borders. **i**. Comparison of Navβ4 expression levels (normalized optical density) across age groups within RA (black) and in an arcopallial region of equal size outside RA (grey). Stars depict significant differences in RA (one-way ANOVA with Tukey’s post hoc; P= 0.005, F (3, 12) = 7.416, N = 4 males per age). **j**. Comparison of the proportions of Navβ4-expressing cells across age groups within RA (black) and in an arcopallial region of equal size outside RA (grey). Stars depict significant age differences in RA (one-way ANOVA with Tukey’s post hoc; P = 0.001, F =F (3, 12) = 10.79, N = 4 males per group). **k**. Comparison of Navβ3 expression level (normalized optical density) across age groups within RA (black) and in an arcopallial region of equal size outside RA (grey). Stars depict significant differences in RA (one-way ANOVA with Tukey’s post hoc; P < 0.0001, F (3, 12) = 73.43, N = 4 males per group). **l**. Comparison of the proportions of Navβ3-expressing cells across age groups within RA (black) and in an arcopallial region of equal size outside RA (grey). Stars depict significant age differences in RA (two-way ANOVA with Tukey’s post hoc; P < 0.0001, F (3, 12) = 90.96, N = 4 males per group). In I-L: * = P ≤ 0.05, ** = P ≤ 0.01, **** = P ≤ 0.0001.

In sharp contrast to RAPNs, neurons recorded in a caudal arcopallial region outside RA (Fig. 2a; shaded area), that has low Navβ4 expression, showed a much smaller I_NaR_ (Fig. 2d-e). Notably, the recorded cells outside RA were morphologically distinct from neurons within RA, with relatively smaller somata and dendrites with numerous, thin spines (Supplementary Fig. 1f). Because I_NaR_ is a function of previously opened Nav channels^52^, the expression of which can vary across cells, we normalized peak I_NaR_ measurements to the peak I_NaT_ (ratio of I_NaR_/I_NaT_). The average normalized peak I_NaR_ was still much larger in RAPNs compared to neurons outside RA over a range of test potentials (−60 mV to −30 mV), with maximum normalized values of 0.28 ± 0.01 and 0.08 ± 0.01 at −45 mV for RAPNs and neurons outside RA, respectively (Fig. 2f). Decay kinetics also showed differences, with RAPNs exhibiting significantly smaller time constants at depolarized test potentials (−15 to +15 mV; Supplementary Fig. 3b).

Because Navβ4 has been linked to facilitation of high frequency firing in mammals and chicken^52,54^, we predicted that RAPNs would be more excitable than neurons outside RA. Indeed, recordings of spontaneous APs proved consistent with this prediction with RAPNs exhibiting narrower APs, larger AP amplitudes and larger maximum rates of depolarization and repolarization than neurons outside RA (Table 1). Furthermore, while neurons both inside and outside RA were capable of increased spiking in response to current injections (Fig. 2g-i), RAPNs produced more spikes per second with a higher instantaneous firing frequency (IFF) for the first two APs at all positive current injections above 200 pA (Fig. 2i-j). These findings point to a strong correlation between Navβ4 mRNA expression, larger I_NaR_, and greater intrinsic excitability, suggesting a role for Navβ4 in regulating intrinsic excitable properties of RAPNs in zebra finches.

### Navβ4 and I_NaR_ increase in parallel in RA during vocal development

Male juvenile zebra finches progress through a critical period of vocal learning, during which vocal practice guided by auditory feedback is required to accurately produce a copy of the tutor song^12,55^. Zebra finches do not sing prior to ∼28 days post hatch (dph). During the next phase (∼28 to 45 dph) they produce unstructured vocalizations, referred to as subsong. They next enter a plastic phase (∼45 to 90 dph), during which songs become more structured as the tutee refines syllable structure and sequencing guided by auditory feedback. By ∼90 dph the song becomes crystallized and highly stereotyped. These developmental changes in song coincide with the growth and maturation of vocal control nuclei in males. In contrast, female finches do not develop song and their song nuclei experience marked reductions in volume, as well as sharp decreases in neuronal cell number and size^56,57^. The morphology, connectivity and electrophysiological properties of RAPNs also change markedly during the song learning period^32,41,42,56-61^.

To investigate whether developmental changes in RA are also associated with changes in the expression of Navβ subunits, we performed *in situ* hybridizations for Navβ4 and Navβ3 in sagittal brain sections from male zebra finches at ages known to be within the pre-song (20 dph), subsong (35 dph), plastic song (50 dph) and crystallized song (>90 dph) stages of vocal development^55,62^. We observed an age-dependent increase in Navβ4 expression level in RA, with significant differences between 20 dph and 35 dph juveniles compared to adults (Fig. 3a-d, and I, black). We also detected an age-dependent increase in the proportion of Navβ4-expressing cells, progressing from 2% of cells relative to Nissl in pre-song juveniles (20 dph) to 24%, 31%, and 63% in 35 dph, 50 dph, and adult birds respectively, with a significant difference between adult and 20 dph finches (Fig. 3j, black). In contrast, no significant changes in Navβ4 mRNA expression, or the proportion of positive cells, were detected outside RA across ages (Fig. 3a-d, and i-j, grey). In stark contrast to Navβ4, we find that Navβ3 expression level and the proportion of Navβ3-expressing cells in RA decreased markedly across ages (Fig. 3e-h, and k-l, black), with little evidence of change outside RA (Fig. 3e-h, and k-l, grey). Finally, we note that while Navβ2 is non-differentially expressed across the arcopallium, Navβ1 expression is higher in RA of adult males compared to the surrounding arcopallium^49^. We therefore asked whether Navβ1 is developmentally regulated. Similar to Navβ4, we find that Navβ1 expression was not present in nucleus RA in 20 dph males, contrasting markedly with the strong Navβ1 expression in adults (Supplementary Fig. 4).

**Figure 4.**
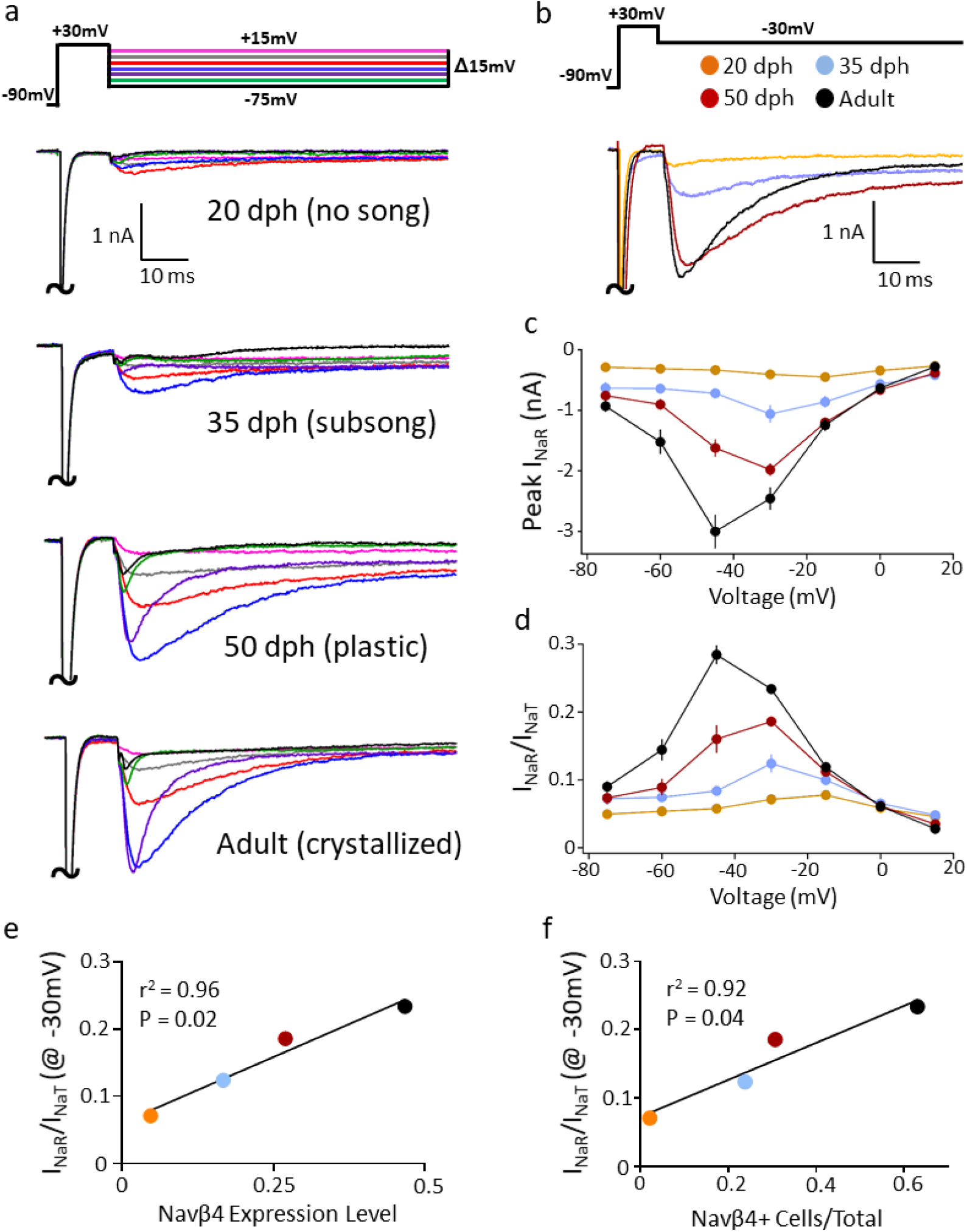
Age-dependent changes in I_NaR_ in RAPNs of male zebra finches. **a**. Examples of I_NaT_ and I_NaR_ elicited across ages by the voltage clamp protocol shown at the top. The large I_NaT_ peaks have been truncated; the color code is the same as in Fig. 2A. **b**. Representative currents from RAPNs at each age group during the −30 mV test potential shown at the top. The large I_NaT_ peaks have been truncated. **c**. Average I-V curves for the peak I_NaR_ in RAPNs from each age group (two-way ANOVA with Tukey’s post hoc comparisons at the −30 mV test pulse; P < 0.0001, F (18, 280) = 16.98, N (cells/age) =10/20 dph, 12/35 dph, 10/50 dph and 12/adults). **d**. Average I-V curves after normalization of the peak I_NaR_ to the peak I_NaT_ measured in a given sweep then averaged across cells (two-way ANOVA with Tukey’s post hoc comparisons at the −30 mV test pulse; P < 0.0001, F (18, 280) = 23.72, N (cells/age) = 10/20 dph, 12/35 dph, 10/50 dph and 12/adults). **e**. Linear regression between average peak I_NaR_ values at the −30 mV test potential across ages and the average Navβ4 expression level in RA. **f**. Linear regression between average peak I_NaR_ values at the −30 mV test potential across ages and the proportion of Navβ4-expressing cells in RA. For views of recorded slices across ages, see Supplementary Figure 5a-c.

Given the changes in Navβ4 expression, we predicted a corresponding age-dependent increase of I_NaR_ in RAPNs. RA could be readily identified in sagittal brain obtained from 20-50 dph male finches via infra-red differential interference microscopy (IR-DIC; Supplementary Fig. 5). Indeed, I_NaR_ was small or absent in RAPNs from 20 dph finches, but its magnitude increased sharply with age (Fig. 4a-b), with significant age-dependent effects observed for both peak I_NaR_ (Fig. 4c) and normalized peak I_NaR_/I_NaT_ ratios (Fig. 4d). At the −30 mV test pulse, significant differences were found for post-hoc pairwise comparisons of raw and normalized I_NaR_ across all age groups. Importantly, the average normalized peak I_NaR_/I_NaT_ values were significantly correlated with both the average Navβ4 expression level (Fig. 4e) and the average proportion of Navβ4-expressing cells within RA (Fig. 4f). Therefore, Navβ4 mRNA expression strongly correlates with I_NaR_ in RA, and predicts the increase of I_NaR_ seen in RA across different ages within the vocal learning period.

**Figure 5.**
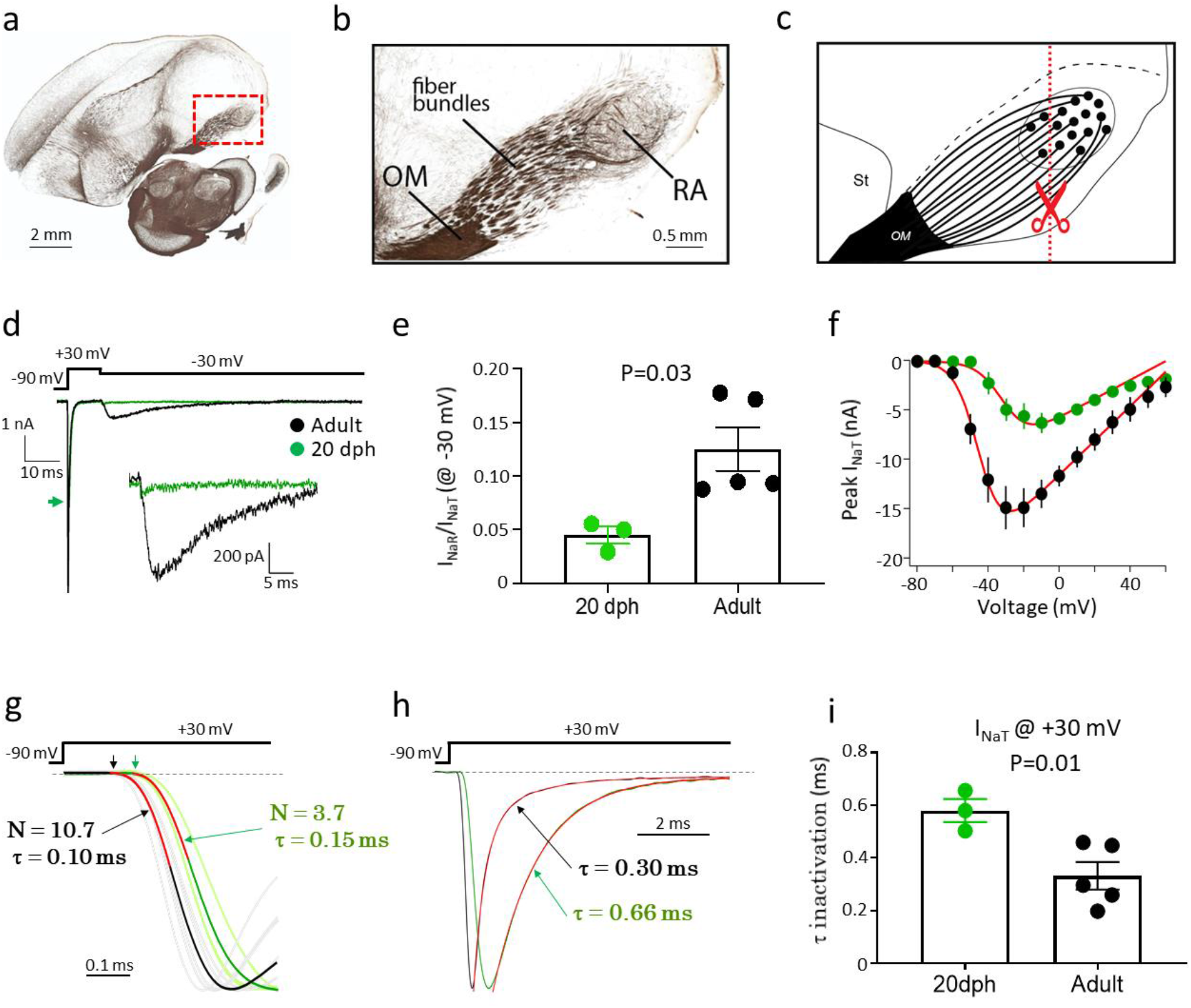
Recording I_NaT_ and I_NaR_ in RAPNs with reduced extracellular Na^+^ in frontal sections from 20 dph and adult male zebra finches. **a**. Myelin-stained parasagittal section containing nucleus RA (from zebrafinchatlas.org^113^; dorsal is up and anterior to the left). **b**. Detailed view of area shown in the dashed red rectangle in (a) depicts myelinated fiber bundles that leave RA, run rostrally, and enter the OM (occipitomesencephalic tract). **c**. Schematic drawing depicts how generating frontal sections (e.g. dashed line with scissors) transect the axonal fibers closer to the somata of RAPNs (black dots in RA), resulting in shorter initial axonal segments of RAPNs than in parasagittal slices. **d**. Overlay of representative currents from 20 dph (green) and adult (black) finches during a −30 mV test potential (top). Green arrow points to the 20 dph I_NaT_ peak. I_NaR_ has been enlarged in the inset. **e**. I_NaR_/I_NaT_ measured at −30 mV in 20 dph and adult finches (N (cells/age) = 3/20 dph and 5/adult finches; Student’s t-test). **f**. Average I-V relationship for I_NaT_ in 20 dph (green) and adult (black) finches. Curve fits for the I-V shown in the red lines. **g**. Curve fit for the current activation shown (red) for averaged I_NaT_ from 20 dph (green) and adult (black) finches where N is the apparent number of transitions in a Markov chain model (equation shown in Methods), τ is the time constant of activation. Scaled I_NaT_ from individual neurons used for averaging shown for adult (grey) and 20 dph (light green) finches. Arrows point to current onsets measured as the first point deviating from a line fit to the pre-stimulus baseline current. **h**. Representative I_NaT_ scaled to the peak from 20 dph and adult; the decay phases have been fitted with single exponentials (red) which overlay the respective traces. **i**. Comparison of the time constant of decay for I_NaT_ in 20 dph and adult finches (N (cells/age) = 3/20 dph and 5/adult males; Student’s t-test).

### Age-dependent changes in the activation and inactivation kinetics of I_NaT_

To address possible concerns that age-dependent differences in our voltage clamp recordings might be affected by space and/or voltage clamp issues, we also performed recordings in frontal slices that transect RA axons near the soma (Fig. 5a-c). For those recordings, we lowered extracellular Na^+^ from 119 to 30 mM to decrease the magnitude of I_Na_ and compensated the series resistance electronically from 5 to 1 MΩ. These changes will improve space and voltage clamp. Consistent with sagittal recordings with normal extracellular Na^+^ (Fig. 4a-d), we observed a significant age-dependent increase in I_NaR_ and confirmed that the normalized peak I_NaR_ was significantly larger in the RAPNs of adults compared to 20 dph finches, where I_NaR_ was almost undetectable (−30 mV test pulse; Fig. 5d-e). We note that we are likely underestimating I_NaR_ in these experiments as lowering external Na^+^ ions disproportionately reduces I_NaR_^52^. Importantly, the resulting I-V plots of the peak I_NaT_ from adults and 20 dph finches suggest that these currents are well voltage-clamped under these conditions (Fig. 5f)^63^. Adults had a larger I_NaT_ amplitude that appeared to activate at more hyperpolarized potentials than that of 20 dph juveniles (Fig. 5f). This likely leads to the faster AP upstroke and decreased threshold in adult RAPNs compared to juveniles (Table 1). In order to quantify the voltage dependence of I_NaT_ we fit the average I-V plots with the following Boltzmann function:

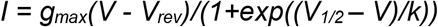

where g_max_ is the maximum conductance, V_rev_ is the reversal potential, V_1/2_ is the half-maximal activation and k is the slope^64^. Fitting this function to the I-V curves revealed that adults had a notably larger maximal conductance and leftward shifted half-maximal activation compared to juveniles. The values from the fits were as follows for adults and juveniles, respectively: g_max_ = 174.0 and 95.0 nS; V_rev_ = 66.5 and 60.0 mV, V_1/2_ = −44.6 and −30.7 mV, and k = 6.6 and 7.3 mV. As predicted by the RAPN firing properties, the averaged I_NaT_ also had a faster onset from the voltage step in adults (0.11 ms) compared to 20 dph juveniles (0.17 ms) (Fig. 5g). In order to quantify the kinetics of I_NaT_ we fit the onset of this current with the following function:

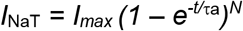

where τ_a_ is the activation time constant and N is the power law exponent for the apparent number of transitions of I_NaT_ activation as defined by a Markov chain model for a classic Hodgkin-Huxley Na^+^ current activation m gate^65^. Interestingly, fitting this function from the onset of I_NaT_ to ∼40% of the peak during a +30 mV test pulse yielded an apparent number of activation gates in the averaged adult I_NaT_ (N = 10.7) that greatly deviated from the Hodgkin-Huxley gating scheme (N = 3); this deviation was less apparent in the juvenile I_NaT_ (N = 3.7; Fig. 5g). Similar fits to I_NaT_ yielded an N = 5 at +30 mV steps from nucleated patches of L5PNs^65^ and a N = 9 at +10 mV steps in cerebellar granule cells^66^. The time constants of activation were also smaller in adults (τ_a_ = 0.10 ms) compared to juveniles (τ_a_ = 0.15 ms) (Fig. 5g). Furthermore, the time constant of inactivation for I_NaT_ was ∼ 2-fold smaller in RAPNs from adults (0.33 ± 0.05 ms; mean ± SE) compared to 20 dph juveniles (0.58 ± 0.04 ms) (Fig. 5h-i). This suggests the presence of a superfast Nav channel inactivation process that precedes the I_NaR_. This fast inactivation may be mediated in part by the fast block provided by Navβ4, as suggested in other systems^51,67^. During these studies we also observed a small persistent Na^+^ current in adult RA (0.3 ± 0.1 nA at 100 ms after a voltage step from −90 to −20 mV at steady state; Fig. 5h).

### Age-dependent increases in the intrinsic excitability of male finch RAPNs

We next investigated the intrinsic excitability of RAPNs during development using whole-cell current clamp recordings. RAPNs showed spontaneous firing at all ages examined (Table 1; representative spontaneous spikes shown in Fig. 6a). The recorded APs exhibited significant age-dependent decreases in threshold and increases in maximum depolarization and repolarization rates (Fig. 6b-c; Table 1). We also found age-dependent changes in the passive properties, with significant decreases in both the input resistance (R_in_), consistent with previous findings in male finches^61^, and in membrane time constant (Table 1). Although the calculated membrane capacitance (C_m_) showed an upward trajectory across ages, we observed a trend toward a decrease from P50 to adults (Fig. 4b; Table 1). This observation mirrors those from a previous study^41^. This decrease in C_m_ may be due to an increase in axonal compact myelination between these ages (Fig.5b and Supplementary Fig. 5;^68^). Furthermore, by multiplying the average C_m_ (114 pF) by the maximum dV/dt we calculated a large peak Na^+^ current of 61 nA in adult RAPNs produced during a spontaneous AP (Table 1). By comparison, neurons of similar size (C_m_ = 80 pF) within deep cerebellar nuclei of mammals have only 21 nA currents^69^. These estimated peak Na^+^ current values increased markedly across age in finches (Table 1), a finding that is supported by voltage clamp experiments (Fig. 5d&f).

**Figure 6.**
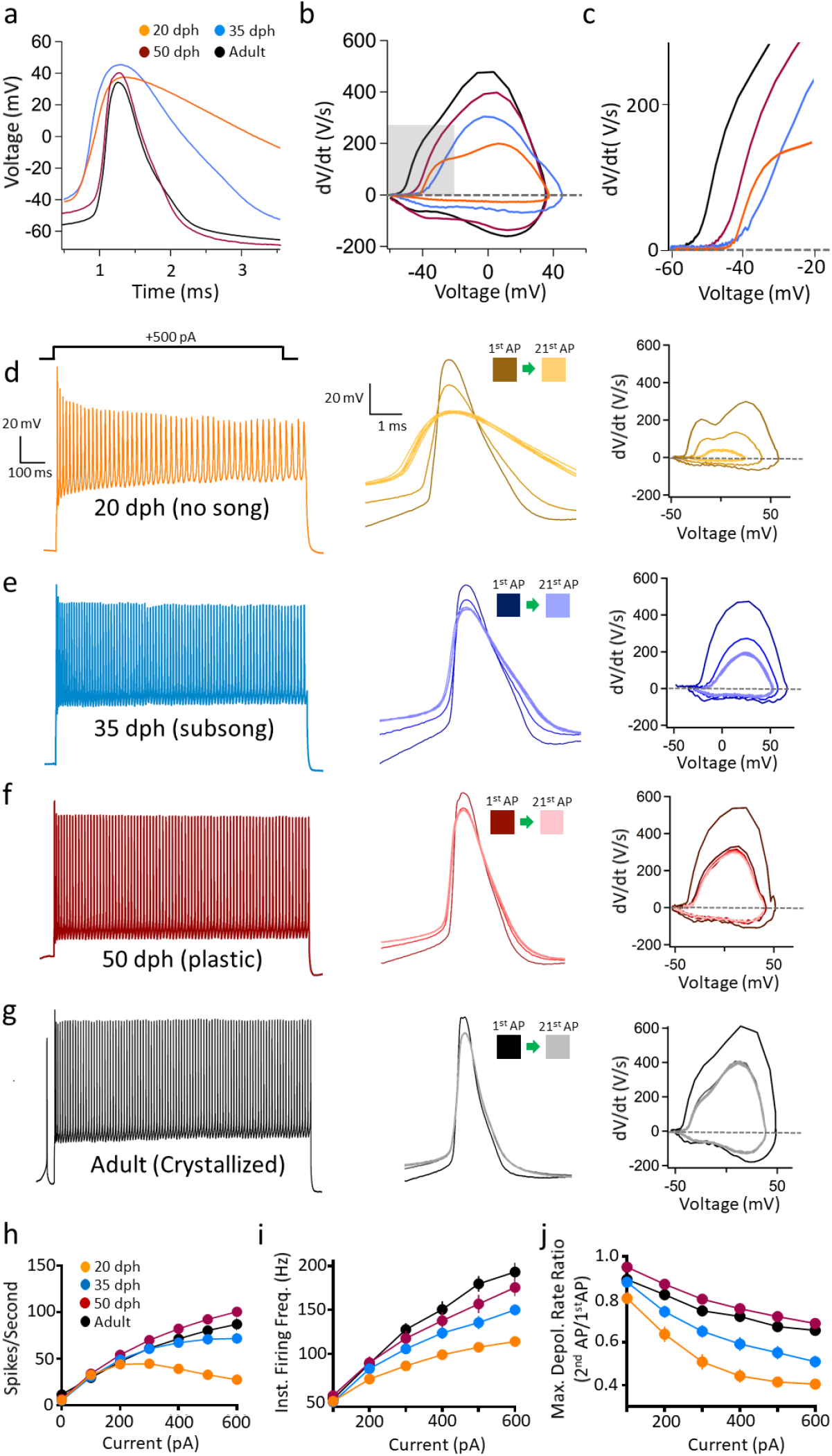
Age-dependent changes in intrinsic excitable properties of RAPNs in male zebra finches. **a**. Overlay of representative spontaneous APs recorded across ages. **b**. Overlay of phase plane plots derived from the APs shown in (a). **c**. Enlarged view of the area highlighted in grey in (b). **d-g**. Left: Representative AP trains elicited by 1 sec 500 pA current injection in RAPNs across ages representative of vocal development stages. Center: APs #1, 2 and 16-21 from the traces on the left aligned at the peaks for each age group. Right: Phase plane plots from APs #1, 2, and 16-21 plotted at the same scale for all age groups. **h**. Average number of spikes produced during 1 sec as a function of current injected for each age group (two-way ANOVA with Tukey’s post hoc; P < 0.0001, F (18, 494) = 4.189, N (cells/age) =18/20 dph, 20/35 dph, 20/50 dph and 18/adults). **i**. Average instantaneous firing frequency as a function of current injected for each age group (two-way ANOVA with Tukey’s post hoc; P = 0.004, F =F (15, 361) = 2.320, N (cells/age) =18/20 dph, 20/35 dph, 20/50 dph and 18/adults). **j**. Fold-change of the maximum depolarization rate from the 1^st^ to the 2^nd^ AP as a function of current injected for each age group (two-way ANOVA with Tukey’s post hoc; P <0.0001, F (15, 348) = 6.774, N (cells/age) =18/20 dph, 20/35 dph, 20/50 dph and 18/adults).

RAPNs at all ages examined were capable of increased AP firing in response to positive current injections (Fig. 6d-g, left). However, we observed a significant age-dependent increase in the number of spikes per second produced during current injections. APs in 20 dph finches began failing at current injections > 200 pA. Significant differences were seen when comparing 35 dph juveniles to either 50 dph or adults at current injections ≥400 pA (Fig. 6h). We also observed a significant age-dependent increase in the IFF, with significant differences when comparing 20 or 35 dph juveniles to either 50 dph or adults at multiple levels of injected current (Fig. 6i). We thus conclude that both spike number and IFF reached a maximum at 50 dph that persisted through adulthood.

We next examined how features of AP waveforms changed during development. By overlaying the 1^st^, 2^nd^ and 16^th^-21^st^ APs elicited by a +500 pA current injection, we were able to observe the effect of repetitive firing on spike amplitude and duration. This analysis revealed a progressive AP broadening and reduction in amplitude in 20 dph birds that began after the 1^st^ AP, eventually reaching a steady state as the train progressed (Fig. 6d-g, middle). These effects were largely attenuated in older juveniles and adults (Fig. 6d-g, middle). Consistent with this finding, phase plane plots of 20 dph APs revealed a marked decrease in maximum depolarization rate with increasing AP number, indicating a loss of Nav channel availability. This effect was noticeably less pronounced in older juveniles and adults (Fig. 6d-g, right). We also found a significant effect of age on the fractional loss in the maximum depolarization rate between the 1^st^ and 2^nd^ AP, with significant differences across all pairwise comparisons during the +600 pA current injection (Fig. 6j). Together, these findings are consistent with the hypothesis that a large I_NaR_ component is important for preserving Nav channel availability during high frequency firing. These findings also provide evidence that the intrinsic excitability of RAPNs undergoes substantial developmental changes, in contrast with previous studies suggesting a lack of marked age-dependent changes^41^.

To facilitate long periods of whole cell recordings we routinely performed these current clamp recordings at room temperature (∼24°C) because recordings did not last long at higher temepratures. However, the gating kinetics of ion channels are highly dependent on temperature, with Q_10_ values varying widely across different ion channels^70^. We therefore performed an additional set of current clamp recordings in 20 dph and adult finch RAPNs at physiological body temperature (∼40°C). Consistent with our room temperature recordings, we found significant differences between 20 dph and adult RA in both spikes number (Supplementary Fig. 6a-c) and IFF (Supplementary Fig. 6d) in response to injected current.

### Navβ4 C-terminal peptide induces I_NaR_ and increases the excitability of juvenile RAPNs

To directly test whether Navβ4 is involved in generating an I_NaR_ in RAPNs, we dialyzed a 14-residue peptide derived from the Navβ4 C-terminus^51^ into RA neurons of 20 dph juveniles, shown previously to possess very low I_NaR_ (Fig. 4a-d; Fig. 5d-e). Consistent with a role for Navβ4 in conferring resurgent properties, intracellular dialysis of the β4-WT peptide elicited a large I_NaR_ within several minutes of achieving the whole-cell voltage clamp configuration (Fig. 7a_1_-a_2_). This effect was not seen with a scrambled version (β4-Scr, 100 μM) of this peptide (Fig. 7c;^51^). The average normalized I_NaR_ to I_NaT_ ratio in RA increased significantly with β4-WT (Fig. 7b) but not with β4-Scr (Fig. 7d) and the fold-change for this ratio differed significantly between cells exposed to β4-WT or β4-Scr (Fig. 7e). We conclude that the channel blocking carboxy-terminal domain of the Navβ4 protein is sufficient to elicit robust I_NaR_ in juvenile RAPNs.

**Figure 7.**
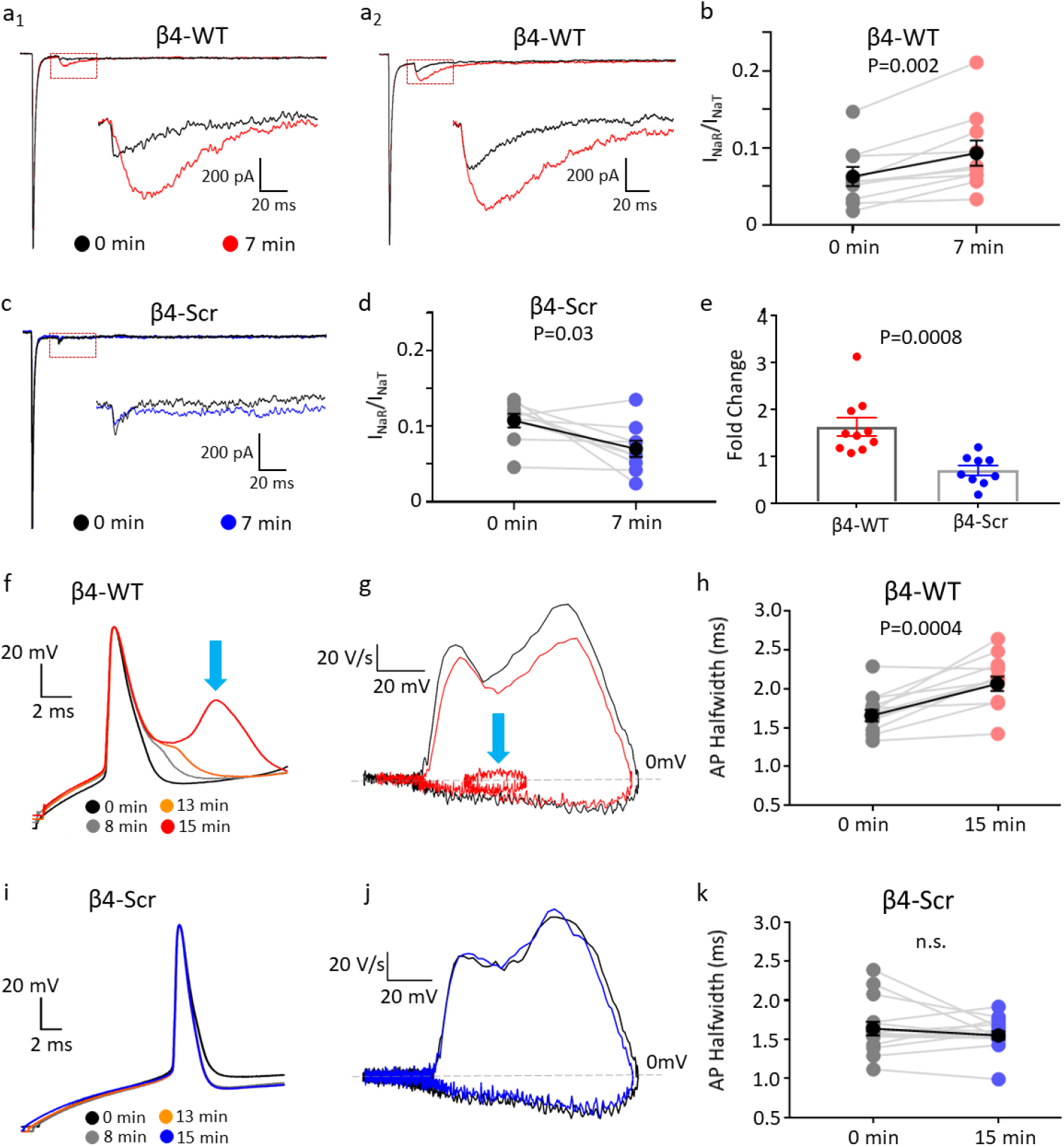
Effects of Navβ4 C-terminal peptide (β4-WT) on I_NaR_ and intrinsic excitability in RAPNs of 20 dph males. **a**_**1-2**_. Representative traces of I_NaT_ elicited at +30 mV followed by I_NaR_ elicited at −30 mV immediately after (black) and 7 min after (red) achieving whole-cell configuration with β4-WT (KKLITFILKKTREK) in the patch pipette. The regions in the red dashed boxes are enlarged in the insets. **b**. Time-dependent increase in the normalized I_NaR_ to I_NaT_ ratio caused by β4-WT, measured at the −30 mV test potential in individual RAPNs (mean ± SEM in black); paired t-test, N= 10 and 9 RAPNs for β4-WT and β4-Scr, respectively. **c**. Representative traces of I_NaT_ elicited at +30 mV followed by I_NaR_ elicited at −30 mV immediately after (black) and 7 min after (blue) achieving whole-cell configuration with β4-Scr (KIKIRFKTKTLELK) in the patch pipette. The region in the red dashed box is enlarged in the inset. **d**. Lack of time-dependent increase in normalized I_NaR_ to I_NaT_ ratio by β4-Scr, measured at the −30 mV test potential in individual RAPNs (mean ± SEM in black); paired t-test, N= 10 and 9 RAPNs for β4-WT and β4-Scr, respectively. **e**. Fold-changes in normalized I_NaR_ to I_NaT_ ratio between 0 and 7 min for RAPNs exposed to β4-WT or β4-Scr in the patch pipette; Student’s t-test; N= 10 and 9 RAPNs for β4-WT and β4-Scr, respectively. **f**. Representative traces of the first APs elicited by a +300 pA current injection at four time points after entering the whole-cell configuration with β4-WT in the patch pipette; the APs are aligned at their peak. **g**. Phase plane plots from the first elicited AP recorded immediately (black) or 15 min (red) after entering the whole-cell configuration in cells exposed to β4-WT. The blue arrows point to the spikelet and resulting alteration in the repolarization rate in the phase plane plot. **h**. Time-dependent effects of β4-WT on the 1st AP half-width in individual cells (mean ± SEM in black; paired t-test). **i**. Representative traces of the first APs elicited by a +300 pA current injection at four time points after entering the whole-cell configuration with β4-Scr in the patch pipette; the APs are aligned at their peak. **j**. Phase plane plots from the first elicited AP recorded immediately (black) or 15 min (red) after entering the whole-cell configuration for cells perfused with the β4-Scr. **k**. Time-dependent effects of β4-Scr on the 1st AP half-width in individual cells (mean ± SEM in black; paired t-test). N= 12 and 15 cells recorded with β4-WT and β4-Scr respectively.

We next examined whether dialyzing 20 dph RAPNs with the β4-WT peptide would make these neurons more excitable. Indeed, upon dialysis of the β4-WT in whole-cell current clamp recordings, we detected a prominent “spikelet” that developed in a time-dependent manner during the repolarization phase of the first AP elicited by a positive current injection (Fig. 7f). This “spikelet” appeared in 58% of cells (7/12 cells) dialyzed with β4-WT, but never in cells dialyzed with the β4-Scr (0/15 cells) (Fig. 7f and i). Exposure to the β4-WT broadened the AP half-width by 20% (Fig. 7h) and decreased the maximum rate of repolarization by an average of 17% (Supplementary Fig. 7a). Importantly, these effects were not seen with the β4-Scr (Fig. 7j-k; Supplementary Fig. 7b). If we treated “spikelets” as APs, the IFF recorded just after the start of dialysis was not different between groups, but after 15 minutes the IFF was significantly higher in cells that received the β4-WT than in cells that received β4-Scr (143 ± 21.2 Hz vs 84.6 ± 3.8 Hz; Mean ± SEM; Supplementary Fig. 7c-d).

### *In silico* modulation of I_NaR_ alters the intrinsic excitability of RAPNs

As a further test of the role of I_NaR_ in the regulation of excitability in RA, we modeled the I_NaR_ from male RAPNs using a previously published model available in the *Senselab* database as a template^71^. Activation of this modeled I_NaR_ is contingent on brief periods of depolarization followed by hyperpolarization of the membrane potential (see Methods). The experimental data (Fig. 8a) matched the modeled I_NaR_ (Fig. 8b) with a close overlap of I-V curves (Fig. 8c) and similar decay kinetics at hyperpolarized potentials (<-20 mV; Fig. 8d).

**Figure 8.**
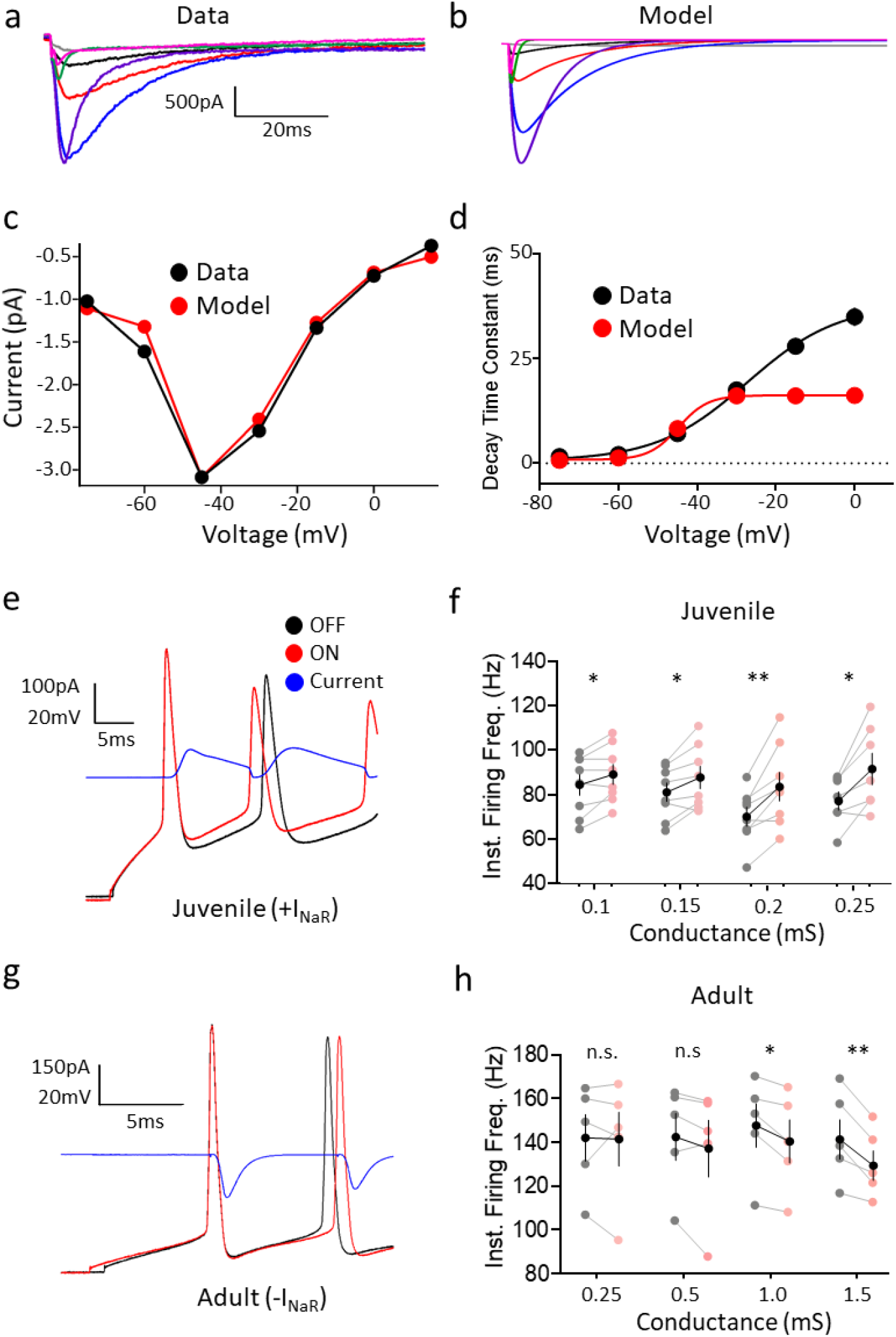
Dynamic clamping of spiking RAPNs in juvenile and adult male finches. **a**. Family of recorded I_NaR_ (same as in Fig. 2C). **b**. Family of modeled I_NaR_ currents (same voltage protocol that elicited currents in Fig. 1c; same scale as in a). **c**. Overlay of the current-voltage relationship of peak I_NaR_ comparing the modeled (red) and average recorded values (black; same as in Fig. 2e). **d**. Overlay of the time constant of decay (τ) as a function of voltage in the modeled (red) and recorded currents (black). **e**. APs elicited in 20 dph RAPNs before (OFF; black) or during dynamic clamp-mediated addition (ON; red) of the *in silico I*_*NaR*_ (blue) peaking during the AP repolarization phase. APs were triggered by a +300 pA current injection. **f**. Paired data of the instantaneous firing frequency change recorded in 20 dph RAPNs upon dynamic clamp-mediated addition (ON; pink) compared to the control (OFF; grey) condition as a function of conductance (mean ± SEM in black, paired t-test). **g**. APs elicited in adult RAPNs before (OFF; black) or during dynamic clamp-mediated subtraction (ON; red) of the *in silico I*_*NaR*_ (blue) peaking during the AP repolarization phase. APs were triggered as in (e). **h**. Paired data of the instantaneous firing frequency change recorded in adult RAPNs upon dynamic clamp-mediated subtraction (ON; pink) compared to the control (OFF; grey) condition as a function of conductance (mean ± SEM in black, paired t-test). N = 8 20 dph and 5 adult RAPNs; In F & G: * = P ≤ 0.05, ** = P ≤ 0.01.

We predicted that adding this *in silico* I_NaR_ via dynamic clamp would enhance aspects of excitability in juvenile RA, while subtracting it would depress excitability in adult RA. We tested these predictions by comparing the effects of delivering sequential +300 pA current injections at baseline and during dynamic clamping of RAPNs. In 20 dph juveniles, we observed that addition of the modeled I_NaR_ during step-like current injections (Fig. 8e, blue) significantly depolarized the interspike periods, while reducing the interval between the first two spikes (Fig. 8e, red; Table 2) when compared to control (Fig. 8e, black; Table 2). This interspike depolarization was expected given the high input resistantance of 20 dph RAPNs (422 MΩ; Table 1). As a result, the IFF increased significantly by 15-20% at all conductances tested (Fig. 8f). These results were reminiscent of our findings using the β4-WT peptide (Supplementary Fig. 7d). This supports a role for I_NaR_ in modulating inter-spike intervals. As with the β4-WT peptide (Fig. 7h and Supplementary Fig.7a), we also noted an increase in the average AP half-width and a decrease in the average maximum repolarization rate (Table 2). Interestingly, the number of elicited APs decreased in a conductance-dependent manner with the injected *in silico* I_NaR_ (Table 2), perhaps due to a low density of voltage-gated K^+^ channels (Kvs) in juveniles that did not allow for a faster recovery of Navs from inactivation.

By contrast, when subtracting the modeled I_NaR_ in adults, we observed hyperpolarization of the membrane potential during the inter-spike periods (Table 3), coinciding with significant decreases in the IFF (5-10%) of the first two spikes when the subtracted *in silico* I_NaR_ conductance was ≥ 1 mS (Fig. 8g-h). Notably, this modulation did not significantly alter the total number of spikes produced during a 1 sec sweep (Table 3). Overall, these findings point to an important role of I_NaR_ in promoting the high-frequency firing of spikes in adult RAPNs.

### Navβ4 mRNA, I_NaR_ and the excitability of RAPNs are sex dependent

Female zebra finches do not sing and show marked developmental atrophy in RA (Supplementary Fig. 5)^56,57^. However, age-dependent changes in the intrinsic excitability of female RAPNs have not been previously examined. *In situ* hybridization for Navβ4 mRNA showed weak expression and no evidence of differential labeling in female RA compared to the surrounding arcopallium at 20 dph, 35 dph or 50 dph (Fig. 9a; refer to Fig. 3 for males). We note that female RA could be unequivocally identified with Nissl staining at the ages tested (Fig. 9b). Contrasting sharply with the marked age-dependent increases of males, we observed a small trend of increased Navβ4 expression after 20 dph (Fig. 9a and c) and a modest, significant increase in the proportion of Navβ4-expressing cells between 20 dph and older ages (Fig 9a and d). Interestingly, these changes appeared to plateau at 35 dph (Fig. 9c and d). Based on this expression pattern, we predicted that the magnitude of I_NaR_ in female RA would be small across ages. Indeed, we detected a small I_NaR_ at 20 dph and a modest increase at 35 dph and no further increase at 50 dph (Fig. 9e-g). However, these trends were not statistically significant.

**Figure 9.**
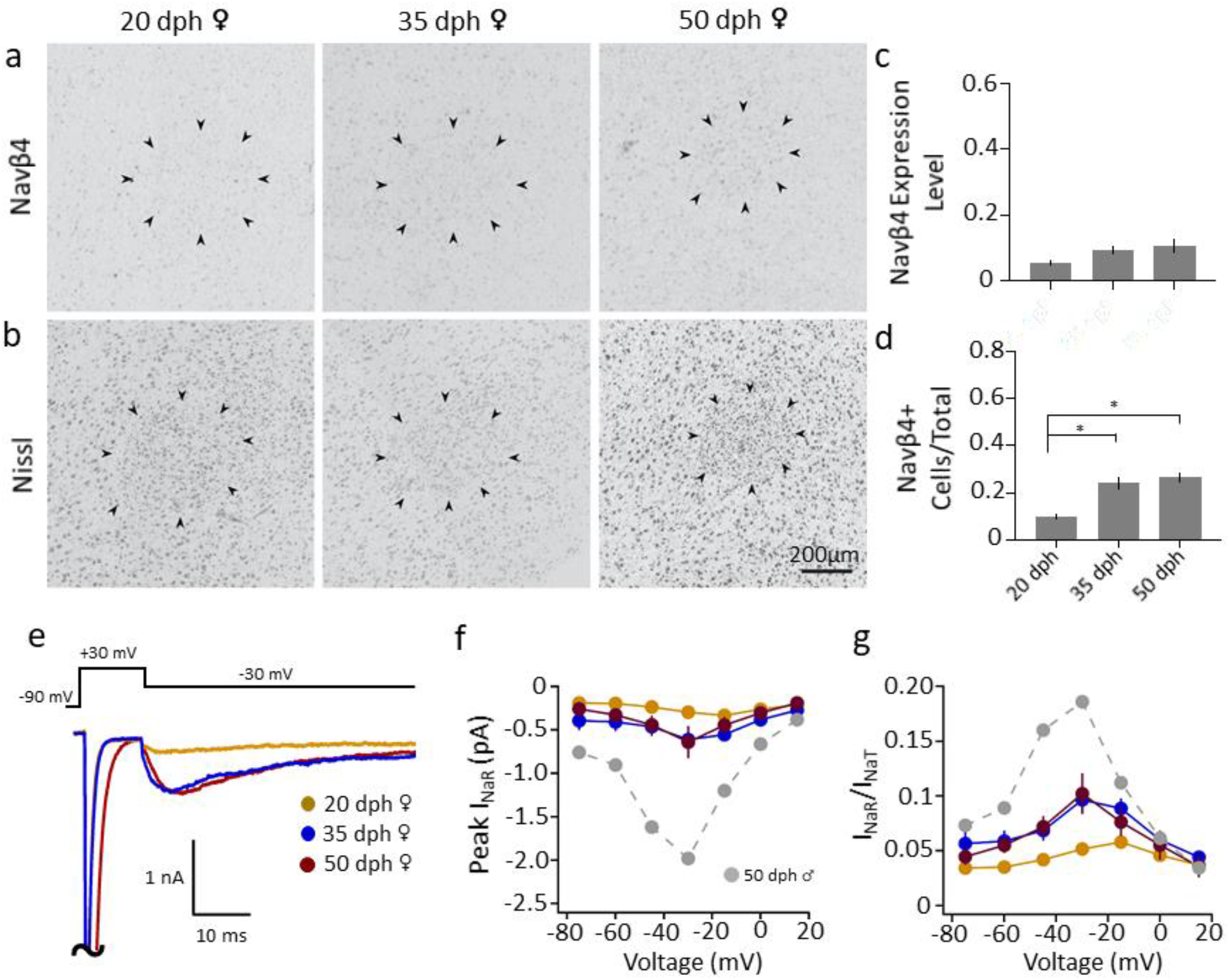
Analysis of Navβ4 mRNA and intrinsic excitable properties in RA of female zebra finches across ages. **a**. Representative *in situ* hybridization images for Navβ4 across ages; black arrowheads indicate RA borders. **b**. Views of adjacent sections to those in (a), stained for Nissl; black arrowheads indicate RA borders. **c**. Comparison of Navβ4 expression levels (normalized optical density) across age groups within RA (one-way ANOVA with Tukey’s post hoc; P = 0.06, 3.290 (2, 10), N (birds/age) = 5/20 dph, 4/35 dph and 4/50 dph females). **d**. Comparison of the proportions of Navβ4-expressing cells across age groups within RA (one-way ANOVA with Tukey’s post hoc; P = 0.0001, F (2, 10) = 24.95; N (birds/age) = 5/20 dph, 4/35dph and 4/50 dph females; stars depict significant age differences in RA determined by post hoc analyses; * = P ≤ 0.01). **e**. Representative currents from RAPNs at each age group during the −30 mV test potential shown at the top. The large I_NaT_ peaks have been truncated. **f**. Average I-V curves for the peak I_NaR_ in RA neurons at each age group; no significant groups differences seen (two-way ANOVA with Tukey’s post hoc; P = 0.83, F (12, 140) = 0.6103, N (cells/age) = 9/20 dph, 10/35 dph and 4/50 dph females). Grey dashed line shows I-V relationship for 50 dph male (same as in Fig. 4c). **g**. I-V curves after normalization of the peak I_NaR_ to the peak I_NaT_ measured in a given sweep then averaged across cells (two-way ANOVA with Tukey’s post hoc; P = 0.48, F (12, 140) = 0.9678, N (cells/age) = 9/20 dph, 10/35 dph and 4/50 dph females). Grey dashed line shows I-V relationship for 50 dph male (same as in Fig. 4d).

Whole-cell current clamp recordings of female RAPNs revealed spontaneous AP firing across all ages, but in contrast to males, there was little evidence of age-dependent changes in active or passive properties (Table 4). These neurons showed increased firing in response to current injections across ages (Fig 10a-c, left, red traces), but unlike males (Fig. 6), we observed a progressive broadening and amplitude decrease when comparing the 1^st^, 2^nd^ and 16^th^-21^st^ AP at all ages examined (Fig. 10a-c, right). We also failed to detect significant age group differences in elicited spikes per sec, in the IFF, or in the change in the maximum depolarization rate ratio between the 2^nd^ and 1^st^ AP (Fig. 10g-i; Supplementary Fig. 8a-c). Phase plots revealed marked differences in rates of depolarization and hyperpolarization between male and female RAPNs at different ages. Specifically, while males had a progressive age-dependent increase in the rate of depolarization during spike trains, the shape of female phase plane plots remained largely unchanged throughout development (Fig. 10d-f). These marked sex differences could be clearly visualized on 3D graphs plotting the relationships between spikes per sec, IFF, and the change in the maximum depolarization rate (Fig. 10g-i). Specifically, while 20 dph males and females were tightly clustered together (Fig. 10g), a moderate separation appeared at 35 dph (Fig. 10h), with the sexes occupying distinct clusters by 50 dph (Fig. 10i). Finally, we note the persistence of the double-hump shape of the female AP phase plot across ages (Fig. 10d-f), similar to that of 20 dph males (Fig. 6b; Fig. 10d, right), and in contrast to 35 dph and 50 dph males. The lack of change in this waveform suggests a persistently low density of Nav channels in the female RAPNs at the axon initial segment and in the soma-dendritic compartment^43^.

**Figure 10.**
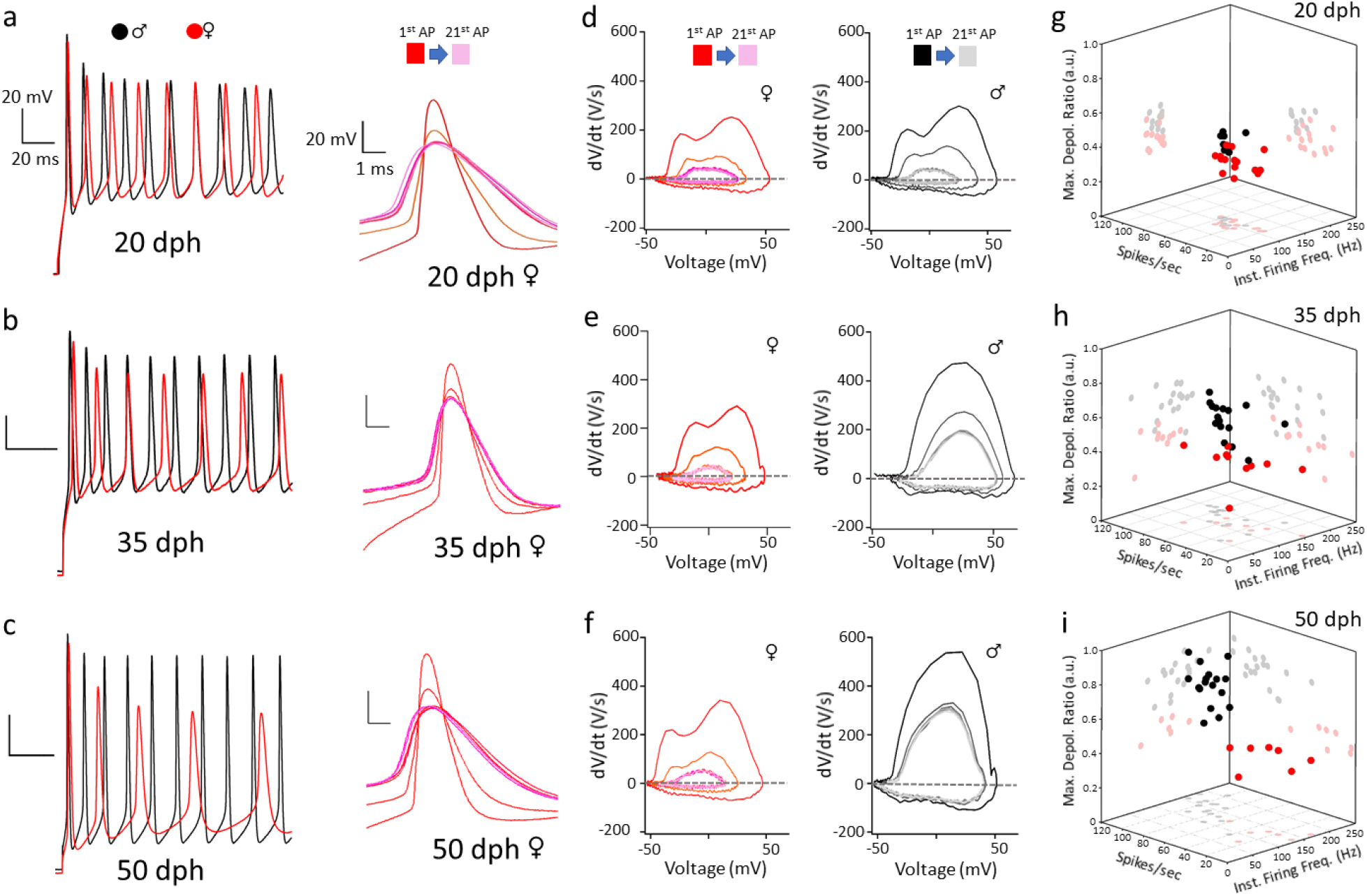
Age-dependent changes in intrinsic excitable properties of RAPNs in female and male zebra finches. **a-c**. Left: Representative AP trains elicited by a 500 pA current injection in RAPNs in males (First 10 APs; black; same as in Fig. 6d-f, left) and females (red) across ages. Right: APs #1, 2 and 16-21 from the traces on the left aligned at the peaks for each age group in females. **d-f**. Phase plane plots from APs #1, 2, and 16-21 plotted at the same scale for all age groups for females (left) and males (right; same as in Fig. 6d-f, right). **g-i**. 3D dot plots showing the instantaneous firing frequency, number of spikes per second and the fold-change in maximum depolarization rate between the 2^nd^ and 1^st^ APs during a 1 sec 500 pA current injection in male (black) and female (red) RAPNs. N (cells/age) = 21/20 dph, 23/35 dph and 7/50 dph females. Refer to Fig. 6 for male data.

Principal component analysis (PCA) applied to multiple spontaneous AP parameters from male and female RA neurons across ages provided quantitative support and further insights into these marked developmental sex differences. PCA1, which accounted for 53% of the total variance, segregated males but not females according to age, with 50 dph males localizing to the right quadrants, whereas younger males and all females from all age groups localized mostly to the left quadrants (Supplementary Fig. 9, x-axis). Interestingly, PCA1 accounted for >80% of the variance of AP half-width and the maximum rates of depolarization and repolarization, half-width showing a strong negative correlation with the other two parameters. PCA2 and PCA3 accounted respectively for 20%, and 17% of the total variance. PCA2 predominantly accounted for the variance of the AP amplitude and threshold, which showed a strong negative correlation, and PCA3 almost exclusively accounted for the variance in the spontaneous firing frequency, but neither PCA2 (Supplementary Fig. 9, y-axis) nor PCA3 (not shown) helped segregate the data by age groups.

## Discussion

Our results reveal that AP waveforms in mature RAPNs have: 1) low spike thresholds, 2) short half-widths, 3) non-adapting amplitudes at high firing frequencies, and 4) large maximum depolarization and repolarization rates (Table 1). These properties strongly resemble those of large motor L5PNs in mammals, and emerge progressively in an age-dependent manner that parallels the known critical period of vocal learning in male, but not female, finches. These changes show a strong correlation with developmental changes in Navβ4 expression and I_NaR_. Experiments utilizing dynamic clamp and Navβ4 peptides in juveniles support the conclusion that I_NaR_ provides an important contribution to the intrinsic excitability of RAPNs. Besides presenting strong evidence linking an I_NaR_ to Navβ4 expression, this work provides insight into the molecular requirements of upper motor neurons that drive spectrally complex, learned vocalizations.

### RAPNs operate as pyramidal-like motor neurons

Precise movements and motor learning in mammals requires the coordinated fast activation and suppression of muscles via output signaling from L5PNs in M1 motor cortex^1^. Likewise, song production in the zebra finch requires fast and precise signaling from RAPNs to activate and suppress syringeal musculature^31^. Despite circuit and gene expression evidence suggesting these neuron types are analogs^16,17,19^, functional evidence at the single neuron biophysical level was previously lacking.

Our study reveals striking similarities between the intrinsic excitability of RAPNs and the “regular spiking” L5PNs found in motor cortex^47,72,73^. During periods of silence, adult RAPNs exhibit periodic spontaneous firing^24,39,40^ that mirrors that of rat L5PNs in M1^72,73^. Like L5PNs^25^, RAPNs have an I_h_-mediated sag^40^ and a persistent sodium current that likely contributes to this spontaneous periodic spiking in both cell types. These currents, together with I_NaR_, may facilitate the depolarizing after potential that follows the AP spike, thus setting the periodicity of the spontaneous spiking (Figure 1c). An additional hallmark of L5PNs is an AP phase plot with a biphasic upstroke phase showing a distinct axon initial segment component with an abrupt “kink” followed by a soma-dendritic component^43^. This closely mirrors our results in Fig. 1e. Voltage-clamp recordings from nucleated patches of L5PNs also reveal a fast non-linear activation of I_NaT_ with a short onset delay^65^. Moreover, the developmental profile of RAPNs closely resembles those of maturing rat L5PNs, including narrowing of spike waveforms and increases in spike firing frequency^26^.

### Narrow action potential spikes may facilitate temporal coding

Pyramidal neurons with multiple dendritic branches and a high density of Na^+^ channels at a distal axonal spike initiation site have decreased effective time constants and exhibit rapid-onset spikes with low thresholds^74-76^. Narrow spikes with rapid onset allow phase-locking to high input stimulation frequencies. Accordingly, large human cortical neurons encode higher bandwidths of synaptic input information with low-threshold, rapid-onset spikes compared to smaller mouse cortical neurons that have higher thresholds and slower spike initiation^77^. Our results suggest that a large I_NaT_ generates the rapid upstroke of the narrow AP in the adult RAPNs, making them well “engineered” to integrate and encode bursts of high-frequency synaptic inputs at their multiple spiny dendrites^24^.

Interestingly, L5PNs in the M1 cortex of awake macaques have large-caliber axons and exhibit narrow AP spikes (0.26 ms)^9^ with fast conduction velocities (∼80 to 90 m/s)^10^. Narrow spikes (half-width = 0.42 ms) are also present in large L5PNs in cat motor cortex^11^. Our results demonstrate a similarly fast activation of I_NaT_ in RAPNs that produces a rapid AP upstroke (Fig. 5g), coupled to a fast I_NaT_ inactivation that helps to produce a narrow AP half-width (0.18 ms at 40°C; Sup. Fig. 2a-c). We also note that while RA neurons in the finch^78^ and Betz-type L5PNs in primate cortex^10^ express Kv3.1, expression of this channel is notably absent in rodent L5PNs^10^. The fact that Kv3.1 currents are associated with fast spiking neurons with narrow APs in several brain areas^79^ may partially explain differences in the spike properties of upper motor neurons in different species. Finally, while our data indicates narrower AP spikes in RA than those previously reported^24,39,40^, this may be due to previous studies using lower temperatures, neurobiotin cell fills^80^, and/or higher external divalent ion concentrations^81^. Taken together, our results thus reveal specialized intrinsic properties of RAPNs that contrast starkly with those in pre-motor HVC neurons^82^ and make them well suited to produce a precise temporal code for fine motor control^31^.

### Fast I_NaT_ inactivation and energy efficient spikes

Prolonged burst firing of non-adapting spikes at high frequencies greatly increases the bandwidth of information processing for RAPNs during song production. This requires a constant operation of Na^+^/K^+^-ATPase pumps, which is energetically expensive. Thus, it is not surprising that RA exhibits dense cytochrome C staining^34^. However, energy demands can be kept within reasonable bounds if the narrow spikes of RAPNs are energy efficient. For example, L5PNs exhibit energy-efficient narrow spikes with reliable spike initiation^83^ and smaller amplitudes at 37°C^46^. Fast I_NaT_ inactivation kinetics is key for energy efficiency because it allows for non-overlapping Na^+^ and K^+^ currents during the upstroke and downstroke of the AP^84-87^. The fast I_NaT_ inactivation of adult RAPNs that we report here (Fig. 5) may thus help to reduce energy consumption. We also note that unlike males, the comparatively smaller female RA, with PNs that fail to fire at high frequencies, display less cytochrome C staining^34^.

### Deveopmental changes in RAPN intrinsic excitability

Previous studies have attributed major *in vivo* changes in RA activity during the vocal learning period primarily to changes in the synaptic strength of inputs from HVC and LMAN^32,41,81,88^. However, we observed significant changes between 35 and 50 dph juveniles in several intrinsic excitability features of RAPNs, including input resistance, spike halfwidth, maximum depolarization and repolarization rates, and spike number during high frequency firing (Fig. 6 and Table 1). Furthermore, while the spike frequency during positive current injections reportedly did not change between 50 dph juveniles and adults^41^, we found significant changes in spike threshold and waveform between these ages. We therefore conclude that in addition to changes in synaptic plasticity, significant changes to intrinsic neuronal excitability may also contribute to the developmental increase in burst firing and decrease in spike variability of RAPNs. Importantly, we also show strong developmental upregulation of both Navβ4 and Navβ1 in addition to downregulation of Navβ3 in RA during the critical period of vocal learning. This change in subunit expression strongly correlated with the appearance of a larger I_NaT_ and I_NaR_, leftward shifts in I_NaT_ activation, and faster I_NaT_ inactivation, all properties thought to facilitate high frequency AP firing. These observations support the notion that developmental changes in RAPN excitability might be linked to changes in modulatory β subunit expression.

### Auxiliary Navβ subunits help to shape spike waveforms in RAPNs

Adult male finches show higher expression of Nav1.6 (SCN8A) in RA compared to the surrounding arcopallium^49^. Importantly, this channel carries the majority of I_NaR_ in a number of cell types^52^. We suggest that a combination of fast inactivation mediated by the Nav1.6 α subunit (hinged-lid mechanism), together with the blocking peptide of the covalently bound Navβ4 may greatly accelerate inactivation kinetics seen in Figure 5h. In addition, the Navβ4 blocking peptide is thought to protect Nav channels from entering a long-lived inactivation state^52^. A large surplus of Nav channels available for opening is critical for limiting cumulative inactivation, thus promoting reliable, high frequency AP firing with little amplitude attenuation^89^. Interestingly, coexpression of Nav1.6 and Navβ4 in HEK cells causes a hyperpolarizing shift in activation (∼8 mV)^90^ similar to what we observed (Fig. 5f; a ∼10 mV hyperpolarizing shift of the Na^+^ current IV curve from 20 dph to adult). Without a high density of available Nav channels, RAPNs would exhibit smaller spike amplitudes, prolonged interspike intervals and AP failures during high frequency firing^89^ as seen in 20 dph juveniles (Fig. 6) and in arcopallial neurons outside RA in adults (Fig. 2).

Adult RA also shows high expression of Navβ1, which bind Nav α subunit non-covalently^91^. In mammals Navβ1 promotes the localization of Nav1.6 to the axon initial segment of L5PNs^92^, and contributes to I_NaR_ in cerebellar granule cells^93^. Nav1.6 exhibits fast inactivation^94,95^ and co-expression with Navβ1 greatly accelerates recovery from inactivation, as well as expression of a small persistent current (∼5% of peak current at +20 mV)^95^. It thus seems likely that high expression of Nav1.6, combined with Navβ1 and Navβ4 may contribute to generating large Na^+^ currents with fast activation and inactivation kinetics in RAPNs^90,94^. Of note, Navβ2 has not been linked to I_NaR_ and is not differentially expressed across the arcopallium^49^.

### Navβ4 peptide and I_NaR_ promote excitability in RAPNs

Although Navβ4 has been linked to I_NaR_ and intrinsic excitability in a number of cell-types^51,67,96,97^, such a link has also been disputed in recent studies where elimination of the Navβ4 gene failed to eliminate this current in cerebellar purkinje neurons^98,99^. However, the knockout was constitutive, and alternative compensatory mechanisms may have enabled this I_NaR_ current^99^. In our study, the β4-WT peptide was sufficient to increase I_NaR_ and the IFF of RAPNs from pre-song juvenile finches in a time dependent manner, providing strong evidence of a causal link between Navβ4 and these excitable properties. Most recordings resulted in a second ‘spikelet’ before the first AP had fully repolarized, in some cases resembling the after-depolarization seen in adult RAPNs (Fig. 1c). A similar effect is also seen at the rodent calyx of Held when this peptide is dialyzed into immature nerve terminals^67^. Fast AP repolarization rates are absent in 20 dph finches (Table 1), which may explain why we did not see a time dependent increase in the number of spikes in these peptide experiments. We suggest that with a faster AP repolarization rate, these “spikelets” would have produced full blown spikes that would have more closely resembled the mature AP waveform and firing.

The dynamic clamp approach we used to further study the role of I_NaR_ in RAPN spiking revealed that the addition of an *in silico* I_NaR_ increases the instantaneous firing frequency of 20 dph RAPNs. Conversely, subtraction of I_NaR_ significantly decreased the instantaneous firing frequency of adult RAPNs (Fig. 8). Because the addition or subtraction of a modeled I_NaR_ has no direct effect on Nav channel availability, like a pharmacological peptide blocker would, these results provide strong evidence that I_NaR_ alone is sufficient to enhance the firing frequency of RAPNs.

Large I_NaR_ and fast I_NaT_ inactivation kinetics are also observed in the mature calyx of Held nerve terminal^67,100^, which exhibits narrow AP spikes that can fire at high frequencies (up to 1 kHz;^101^). Moreover, I_NaR_ generates a depolarizing after potential at the calyx of Held once the spike downstroke has terminated^67^, similar to that seen in RAPN spikes (Figure 1c). Interestingly, cells in layer 5 of mouse primary motor cortex (M1) also express Navβ4 mRNA^102^ and Navβ4 protein is specifically expressed at the axon initial segment of L5PNs of the mouse somatosensory cortex^103^, although it is currently unknown whether L5PNs have I_NaR_.

### Sex differences in the intrinsic excitability of maturing RAPNs

Although largely unexplored, defining sex differences in the excitability of zebra finch song nuclei may help to identify critical requirements for neurons that control learned vocalizations. Interestingly, the developmental changes in Navβ4 expression, I_NaR_, spike waveforms and firing rates of RAPNs seen in males were absent in females, which do not sing (Figure 10). These observations provide further support for Navβ4 expression underlying I_NaR_, and thus contributing to the intrinsic excitability properties of RAPNs. While we acknowledge that a number of other factors likely determine the lack of female vocal learning^56,104^, we suggest that the lack of maturation in the intrinsic properties of RAPNs could contribute to females being unable to produce complex, learned vocalizations.

## Conclusions

Our results point to a key piece of “hardware” that allows adult male RAPNs to function as precise and reliable generators of fast spiking: High expression of Navβ4, likely coupled with Navβ1 and Nav1.6α subunits, to produce a large I_NaR_ and an I_NaT_ with fast activation and inactivation. These features lower AP threshold and allow robust rapid-onset APs with large non-adapting amplitudes^76^. Possibly in concert with Kv3.1 channels^10,79^, this combination of Navα/β subunits narrows the spike half-width and enables fail-safe fast spiking activity to emerge during the critical song learning period. By contrast, RAPNs in female and male juveniles express high levels of Navβ3, do not have a large I_NaR_, and cannot spike at high frequencies. Finally, we note that a subclass of large Betz-type L5PNs in primate motor cortex^9,10^ also exhibit narrow spikes that may fine tune them for rapid and precise motor control^3^. We thus propose that the specialized “hardware” of RAPNs facilitate their precise *in vivo* spike burst firing that is required for activation of lower motor neurons and contraction of the superfast syrinx muscles involved in singing^33^.

## Methods

### Animal subjects

All of the work described in this study was approved OHSU’s Institutional Animal Care and Use Committee and is in accordance with NIH guidelines. Zebra finches (*Taeniopygia guttata*) were obtained from our own breeding colony or purchased from local breeders. The ages of birds in the 20, 35, and 50 days post hatch (dph) groups were established based on the date the first egg hatched, and thus the birds were sacrificed within ± 2 days of the target age. Birds older than 120 dph were considered adults. On the day of the experiment, all of the birds with the exception of 20 dph birds, were removed from the colony at lights-on (9 AM PST) and housed for ∼1 hr in an acoustic isolation chamber to minimize the potential confounds of singing and non-specific auditory stimulation on gene expression, noting that social isolation can also alter neurogenomic state^105^. The 20 dph finches were removed directly from the colony, and not isolated, so as to minimize stress on newly fledged birds. The sex of 20 and 35 dph birds could usually be distinguished by plumage, however we routinely confirmed the sex of individuals by gonadal inspection. Birds were sacrificed by decapitation and their brains removed. For electrophysiology experiments brains were bisected along the midline, immersed in ice-cold cutting solution, and processed as described below. For a subset of these brains (N=4 per sex per age group), we reserved one hemisphere for electrophysiology experiments and the other was placed in a plastic mold, covered with ice-cold Tissue-Tek OCT (Sakura-Finetek; Torrance, CA), and frozen in a dry ice/isopropanol slurry for *in situ* hybridization. The use of right vs. left hemispheres was balanced to account for any hemispheric differences in electrophysiology or gene expression. For the remaining brains, both sides were used for electrophysiological recordings.

### *In situ* hybridization

To compare mRNA expression levels for *SCN3B* and *SCN4B* across developmental ages and between sexes, brains sections (thickness = 10 μm) were cut on a cryostat onto glass microscope slides (Superfrost plus; Fisher Scientific) and stored at −80°C. For each brain, a set of slides consisting of every 10^th^ slide was fixed and stained for Nissl using an established protocol. Slides were examined under dark- and bright-field microscopy to identify sections containing the core region of nucleus RA, i.e. the sections where RA appears largest. *In situ* hybridizations (ISH) were conducted using established protocols^106^ and were performed in separate batches for each gene to standardize fixation and hybridization conditions. Briefly, slides were hybridized under pre-optimized conditions with DIG-labeled riboprobes synthesized from BSSHII digested cDNA clones obtained from the ESTIMA: Songbird clone collection^107^, corresponding to GenBank entries FE734016 (SCN3B; aka Navβ3), FE730991 (SCN4B; Navβ4), and DV957065 (SCN1B; Navβ1). After hybridization, slides were washed, blocked, incubated with alkaline phosphatase conjugated anti-DIG antibody (Roche, 1:600), and developed overnight in BCIP/NBT chromagen (Perkin Elmer). Slides were coverslipped with VectaMount (Vector) permanent mounting medium, and then digitally photographed at 10X under brightfield illumination with a Lumina HR camera mounted on a Nikon E600 microscope using standardized filter and camera settings. Images were stored as Tiff files and analyzed further using the FIJI distribution of ImageJ^108^. We note that high-resolution images of SCN1B, SCN3B, and SCN4B expression in the adult male zebra finch brain are available on the Zebra Finch Expression Brain Expression Atlas (ZEBrA; www.zebrafinchatlas.org).

### Image Analysis and Expression Quantification

For each image, we quantified both expression levels based on labeling intensity (i.e. average pixel intensity) and the number of cells expressing mRNA per unit area. Since the density of cells in RA is known to change during development and to vary by sex^56^, we first estimated the number of neurons in RA and caudal to RA for each bird by placing a 200 × 200 μm window over target areas in the images of the Nissl-stained sections containing the largest RA per bird. We next measured the average pixel intensity (scale: 0-256) in an identical 200 × 200 µm window placed over each target area in the images of hybridized sections adjacent to the Nissl-stained sections. From the values we then subtracted an average background level measured over an adjacent control area in the intermediate arcopallium that was deemed to have no mRNA expression. Finally, we divided the background corrected pixel intensity value by the number of Nissl-counted cells in order to obtain a measurement of the average pixel intensity per cell. We also quantified the number of labeled cells in each arcopallial region by first establishing a threshold of expression 2.5X above the background level. Standard binary filters were applied and the FIJI ‘Analyze Particles’ algorithm was used to count the number of labeled cells per 200 µm^2^. This value was further normalized for comparisons across different ages and sex by dividing by the number of Nissl-labeled cells from the adjacent section.

### Slice Preparation for Electrophysiology Experiments

Sagittal (200 μm thick) and frontal (150 μm thick) slices were cut on a vibratome slicer (VT1000, Leica) in an ice-cold cutting solution containing (in mM): 119 NaCl, 2.5 KCl, 8 MgSO_4_, 16.2 NaHCO_3_, 10 HEPES, 1 NaH_2_PO_4_, 0.5 CaCl_2_, 11 D-Glucose, 35 Sucrose pH 7.3-7.4 when bubbled with carbogen (95% O_2_, 5% CO_2_; osmolarity ∼330-340 mOsm). Slices were then transferred to an incubation chamber containing artificial cerebral spinal fluid (aCSF) with (in mM): 119 NaCl, 2.5 KCl, 1.3 MgSO_4_, 26.2 NaHCO_3_, 1 NaH_2_PO_4_, 1.5 CaCl_2_, 11 D-Glucose, 35 Sucrose pH 7.3-7.4 when bubbled with carbogen (95% O_2_, 5% CO_2_; osmolarity ∼330-340 mOsm) for 10 min at 37°C, followed by a room temperature incubation for ∼30 min prior to start of electrophysiology experiments. For low-sodium aCSF experiments (see Fig. 5), 119 NaCl mM extracellular sodium was replaced with 30 mM NaCl and 90 mM NMDG, the pH was adjusted to 7.4 with HCl, and the osmolarity was adjusted to ∼330-340 with sucrose.

### Patch Clamp Electrophysiology

RA could be readily visualized in males and females of all ages via infra-red differential interference contrast microscopy (IR-DIC) (Supplementary Fig. 5). Whole-cell patch-clamp recordings were performed at room temperature (∼24°C) unless otherwise indicated. For experiments performed at 40°C, the bath solution was warmed using an in-line heater (Warner Instruments, Hamden, CT). The temperature for these experiments varied up to ± 2°C. Slices were perfused with carbogen-bubbled aCSF (1-2ml/min) and neurons were visualized with a IR-DIC microscope (Zeiss Examiner.A1) under a 40x water immersion lens coupled to a CCD camera (Q-Click; Q-imaging, Surrey, BC, Canada). Whole-cell voltage- and current-clamp recordings were made using a HEKA EPC-10/2 amplifier controlled by Patchmaster software (HEKA, Ludwigshafen/Rhein, Germany). Data were acquired at 40 kHz and low-pass filtered at 2.9 kHz. Patch pipettes were pulled from standard borosilicate capillary glass (WPI, Sarasota, FL) with a P97 puller (Sutter Instruments, Novato, CA). All recording pipettes had a 3.0 to 6.0 MΩ open-tip resistance in the bath solution. Electrophysiology data were analyzed off-line using custom written routines in IGOR Pro (WaveMetrics, Lake Oswego, OR).

For voltage clamp (VC) recordings, intracellular solutions contained the following (in mM): 142.5 Cs-Gluconate, 11 CsCl, 5.5 Na_2_-phosphocreatine, 10.9 HEPES, 5.5 EGTA, 10.9 TEA-Cl, 4.2 Mg-ATP, and 0.545 GTP, pH adjusted to 7.3 with CsOH, ∼330-340 mOsm. Alexafluor 488 hydrazide (Thermofisher; 10 μM) was added to this solution to aid in cell-type morphology discrimination via epifluorescent microscopy (X-Cite series 120, Excelitas Technologies, Waltham, MA). In recordings where the series resistance (R_s_) was compensated to 5 MΩ, the average uncompensated R_s_ = 10.2 ± 0.5 MΩ (n=66 cells for male/female). In recordings in Fig. 5 where R_s_ was compensated to 1 MΩ, the uncompensated R_s_ = 7.2 ± 0.5 MΩ (n=8 for males at 20 dph and adults). In order to isolate Na^+^ currents, slices were exposed to bath applied CdCl_2_ (100 μM), 4-AP (100 μM), Picrotoxin (100 μM), TEA (10 mM), CNQX (10 μM), and APV (100 μM) for ∼5 min prior to running voltage clamp protocols. After protocols were applied, tetrodotoxin (TTX 1 *μ*M) was bath applied and the same voltage-clamp protocols were repeated after Na^+^ currents were eliminated. Na^+^ currents were isolated by subtracting the TTX-insensitive current traces from the initial traces. Capacitive currents generated during voltage-clamp recordings were eliminated by P/4 subtraction. Recordings were not corrected for a calculated liquid junction potential of 12 mV.

For current clamp recordings (CC), intracellular solutions contained (in mM): 142.5 K-Gluconate, 21.9 KCl, 5.5 Na_2_-phosphocreatine, 10.9 HEPES, 5.5 EGTA, 4.2 Mg-ATP, 0.545 GTP, and 10 μM Alexfluor 488 hydrazide, pH adjusted to 7.3 with KOH, ∼330-340 mOsm. Synaptic currents were blocked by bath applying Picrotoxin (100 μM), APV (100 μM), and CNQX (10 μM) (Tocris Bioscience) for ∼3 min prior to all recordings. To initiate CC recordings, we first established a gigaohm seal in the VC configuration, set the pipette capacitance compensation (C-fast), and then set the voltage command to −70 mV. We then applied negative pressure to break into the cell. Once stable, we switched to the fast CC configuration. Experiments in CC were carried out within a 15-min period. We noted that the resting membrane potential tended to hyperpolarize to the same degree (∼10 mV) in both juveniles an adults during these CC recordings^109^. Recordings in which the resting membrane potential deviated by > 15 mV were discarded. We note that recordings were not corrected for a calculated liquid junction potential of 9 mV.

Estimated current clamp measurements of membrane capacitance (C_m_) were calculated from the measured membrane time constant (τ _m_) and input resistance (R_in_) using the equation:

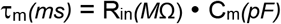

Estimated peak Na^+^ current (I_Na_) activated during the upstroke of the action potential (AP) was calculated from the capacitive membrane current using the equation:

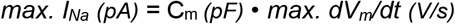

where peak depolarization rate is *max. dV*_*m*_*/dt* (see Vaaga et al. (2020)^69^ and Yu et al. (2012)^46^).

### Dialyzed Peptides

Mouse β4-WT (KKLITFILKKTREK) and β4-Scr (KIKIRFKTKTLELK) peptides (as previously described in Grieco et al. 2005^51^) were synthesized by Thermofisher scientific at 98% purity and diluted to 100 μm in internal solutions for CC and VC experiments. The mouse peptide was used because the C-terminal domain of Navβ4 is highly conserved across vertebrates and found to be readily soluble in our internal solutions^53^.

### Dynamic Clamp parameters

Dynamic clamp was implemented in a Hewlett Packard PC (Z600 workstation) running Windows 10 software^110^. Voltage values were read and calculated currents outputted through a National Instruments DAQ board (PCIe-6321) controlled by Igor Pro 8 using the NIDAQ Tolls MX package (Wavemetrics). We utilized a model deposited in ModelDB within the *Senselab* database (see accession number 127021)^71,111^. The I_NaR_ current was modeled as a single gate relying into two particles: one for activating (*s*) and one for inactivating (*f*). Each of these particles has a separate probability for opening (i.e. going to a permissive state: α) and closing (i.e. going to a non-permissive state: β). In order to match the kinetics of the model to zebra finch I_NaR_ currents, we made minor changes to the equations describing the rate constants in the original model^71^. We adjusted two parameters of the s particle rate constants: (1) an exponential term denominator (Kα_s_ from −6.82 to −1.0) and (2) the magnitude of the multiplying term β rate constant (Aβ_s_ from 0.0156 to 0.05).

To calculate the *in silico* current, we used a set of first order Hodgin-Huxley type equations:

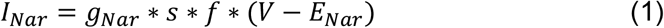

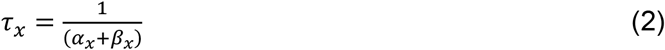

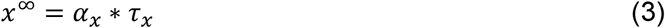

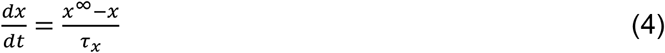

Where *x* represents either the s or f gating particles. The differential equation (4) was solved using a first order Euler method^112^ with a time step of 0.03 ms. The g_Nar_ represents the maximum conductance (in mS) and the calculated reversal potential (E_Nar_) was +67 mV. All neurons recorded were kept approximately at −70 mV throughout the current clamp recordings. To trigger action potentials, a step current injection of +300 pA (relative to the holding current) was applied for 1 sec. A complete list of the numeric parameters and equations is available in ModelDB (modeldb.yale.edu/127021)^71,111^.

### Statistical Data Analysis and Curve Fitting

Data were analyzed off-line using IgorPro software (Wave-metrics). Statistical analyses were performed using Prism 4.0 (GraphPad). Specific statistical tests and outcomes for each analysis performed are indicated in the respective Figure Legends and Tables. Means and SE are reported, unless otherwise noted.

## Acknowledgements

We would like to thank Samatha Friedrich for help with zebra finch breeding. We would also like to thank Pepe Alcami for feedback regarding this manuscript. This work was funded by NIH grants NSF1456302, NSF1645199, NIH/GM120464, NIHDC004274, NIHDC012938.

## Author Contributions

B.M.Z, P.V.L, C.V.M and H.v.G designed the research. B.M.Z, A.A.N., A.D. performed the research, B.M.Z., A.A.N., A.D., P.V.L, C.V.M. and H.v.G. analyzed the data, B.M.Z, P.V.L, C.V.M and H.v.G wrote the paper.

