## Supplemental Figures & Tables for "Resurgent Na^+^ currents promote ultrafast spiking in projection neurons that drive fine motor control"

### Supplemental Figures and Legends:

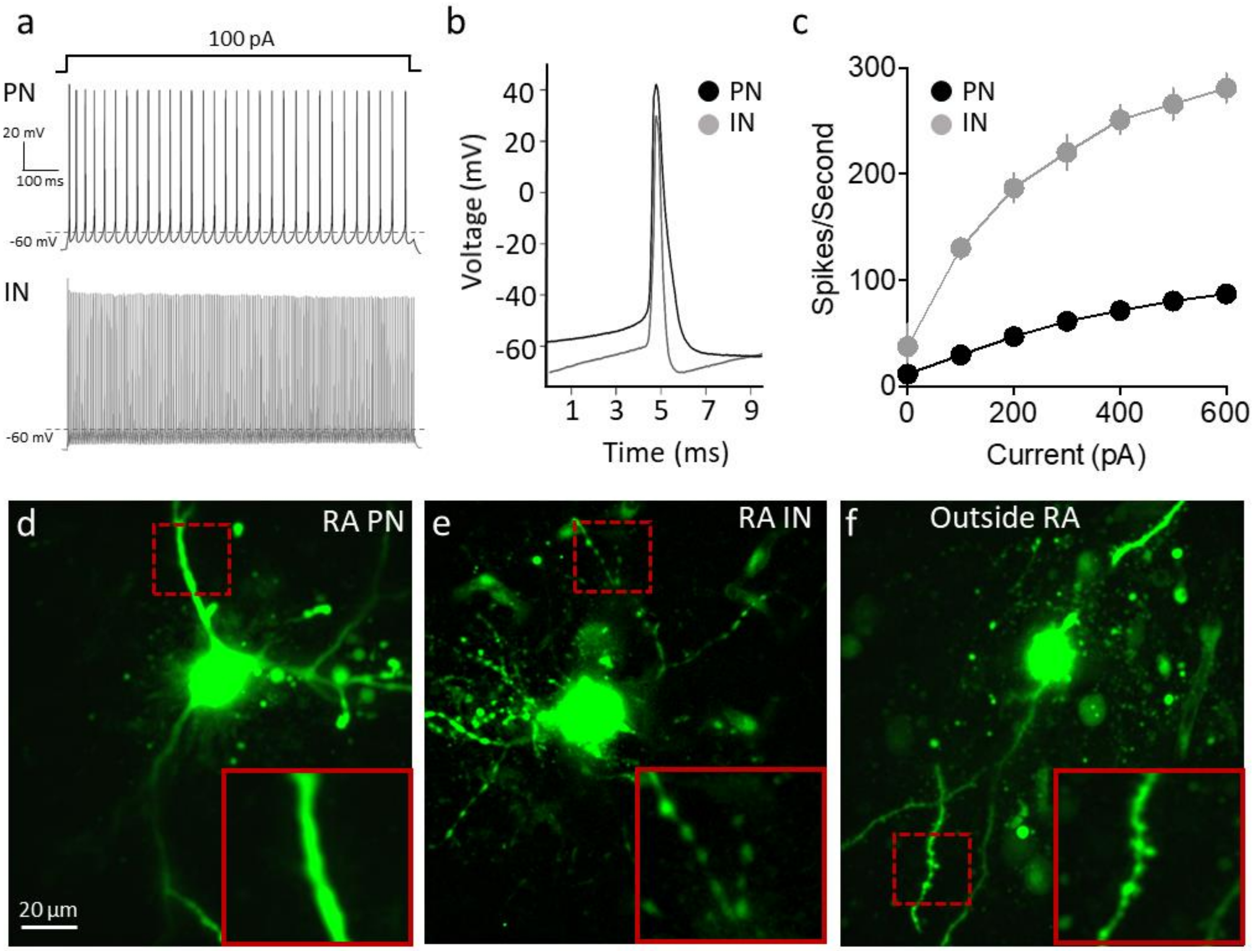


Supplementary Figure 1. Distinct morphology and excitable properties of arcopallial neurons in adult male zebra finches. **a.** AP trains elicited in a RAPN (projection neuron; top) and a RAIN (inhibitory interneuron; bottom) during a 1 sec 100 pA current injection at 24$^{o}$C. **b.** Overlay of average from the first 5 APs in an RAPN and a RAIN (same cells as in A). **c.** Average elicited spikes/sec as a function of current injected in RAPNs and RAINs. **d.** Image of a RA projection neuron (PN) that has been filled with a fluorescent dye (Alexafluor 488 hydrazide) during an electrophysiological recording. Note the lack of varicosities in its processes (dashed box expanded in the bottom right). **e.** Image of an RA interneuron (IN) that has been filled with fluorescent dye as in (d). Note the prominent varicosities in its processes[24,39] (dashed box expanded in the bottom right). **f.** Image of an arcopallial neuron outside RA that has been filled with fluorescent dye as in (d-e). Note the dendritic processes with large spines that are distinct from RAPNs and INs (dashed box expanded in the bottom right).

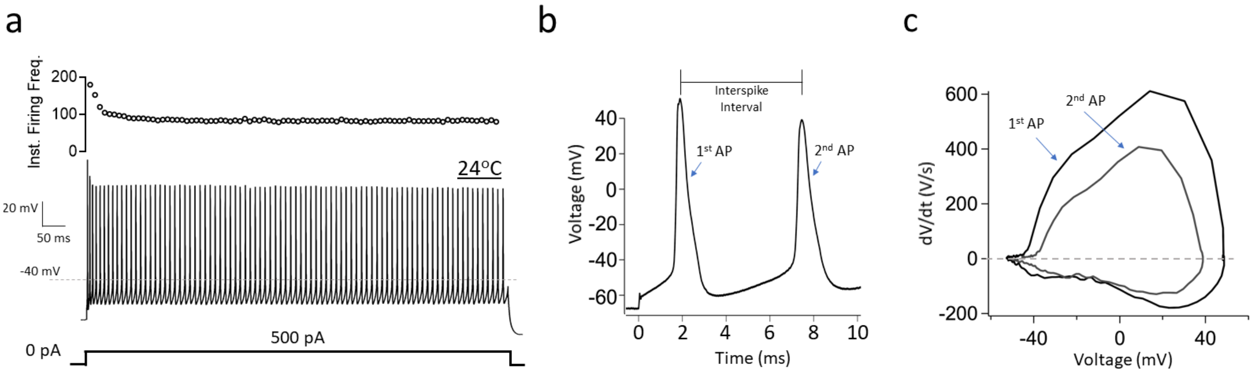


Supplementary Figure 2. Intrinsic excitable properties of RAPNs at room temperature. **a.** Representative AP train (from the same cell as in Fig. 1b) elicited during a 1 sec 500 pA current injection. Top: instantaneous firing frequency plotted against time. **b.** First two APs from (a). **c.** Overlay of the phase plane plots from the two APs in (b).

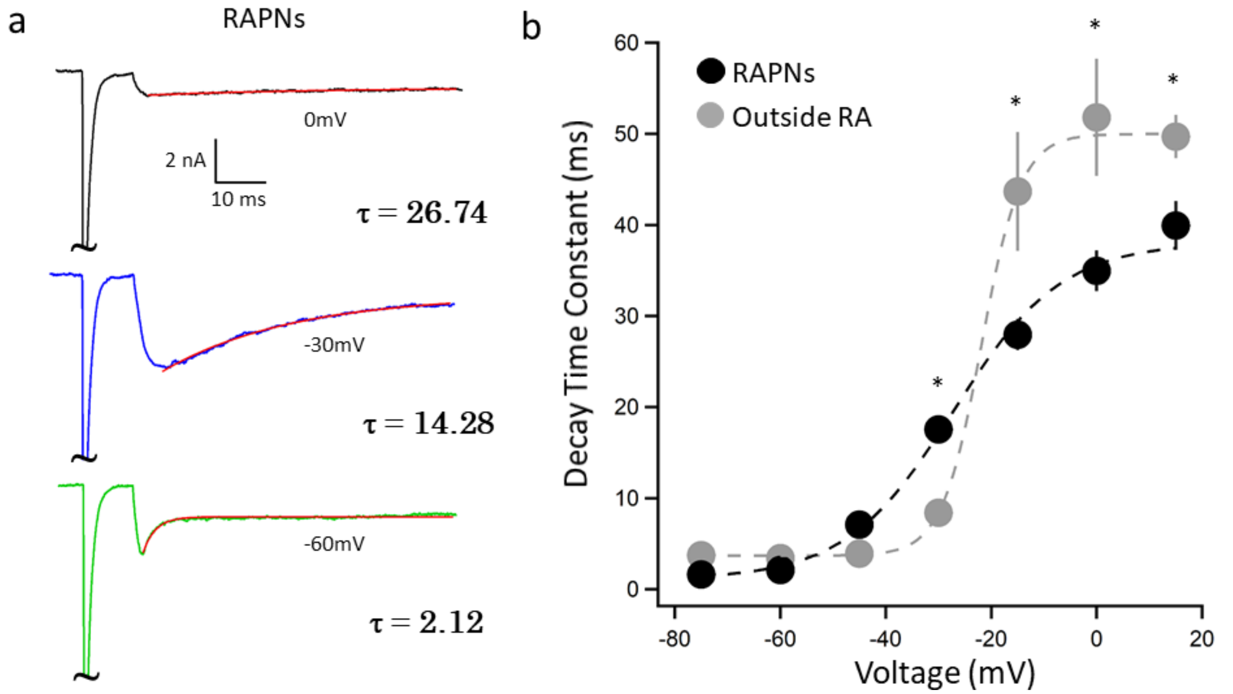


Supplementary Figure 3. Voltage-clamp recordings of arcopallial neurons in slices of adult male zebra finches. **a.** Individual current traces for multiple test potentials overlaid with single exponential decay fits (red trace) for $I_{NaR}$ in RAPNs. $I_{NaT}$ peaks have been truncated. **b.** $I_{NaR}$ decay time constant as a function of test potential for RAPNs and neurons outside RA. Two-way ANOVA with Tukey's post hoc; $P < 0.0001$, $F (6, 126) = 7.628$, N =12 RAPNs and 11 neurons outside RA. Stars depict significant age differences in RA determined by post hoc analyses; * = $P \leq 0.05$.

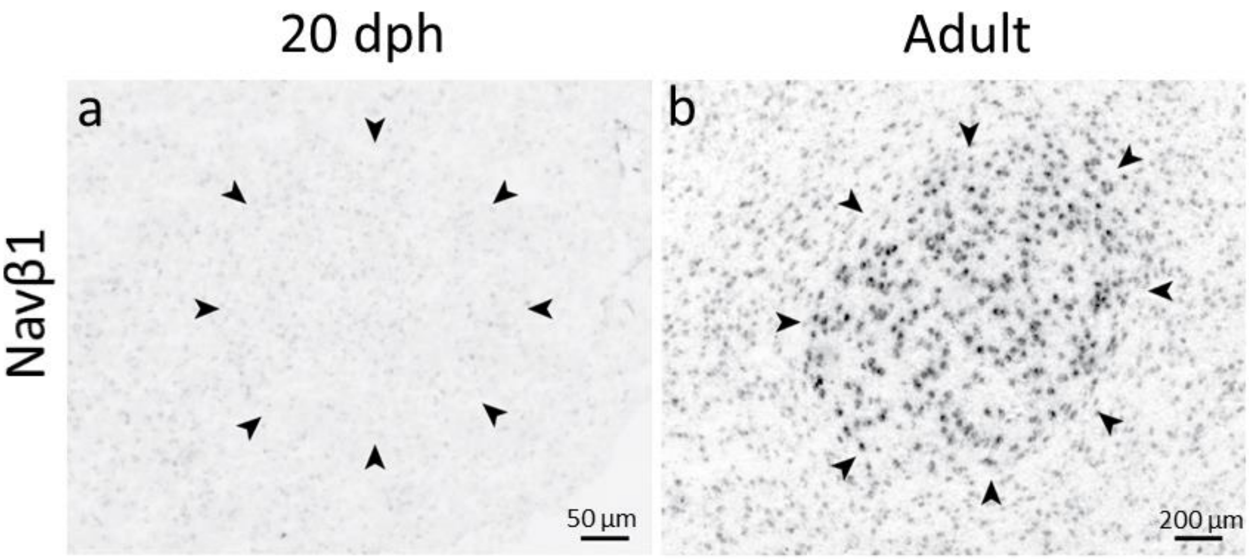


Supplementary Figure 4. Age-dependent change in expression of Navβ1 mRNA in the arcopallium of male zebra finches. **a-b.** Representative *in situ* hybridization images for Navβ1 mRNA in 20 dph and adult zebra finch arcopallium; black arrowheads indicate RA borders.

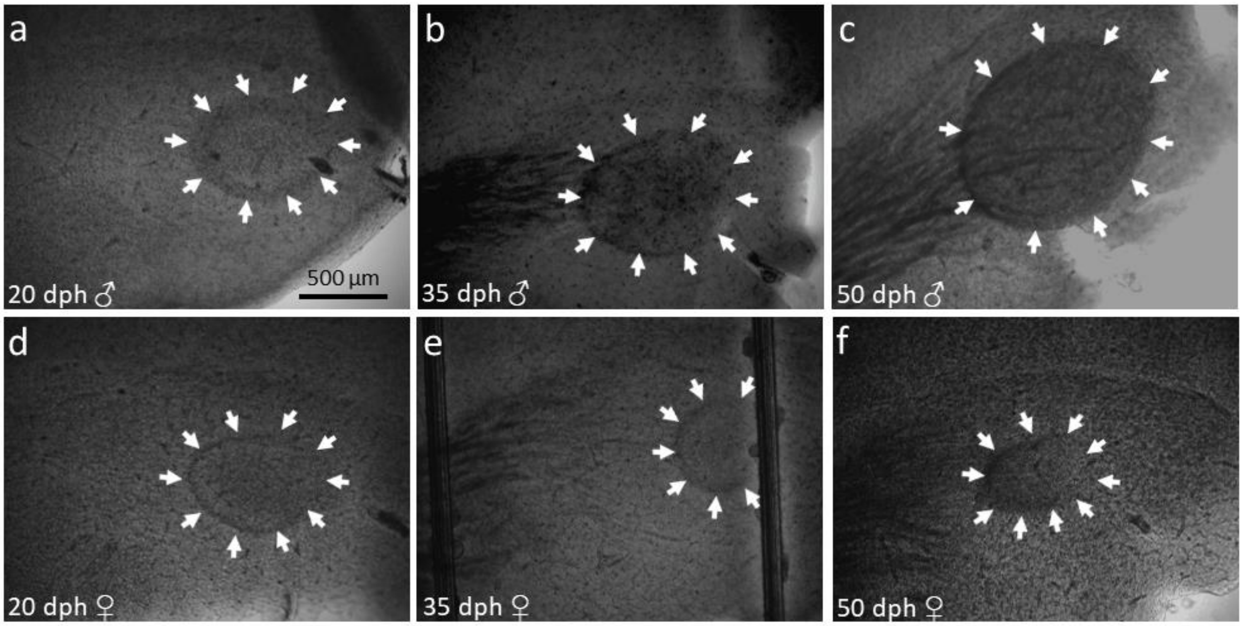


Supplementary Figure 5. Visualization of RA in brain slices from zebra finches across ages. **a-c.** Representative images of parasagittal slices through the arcopallium of 20, 35, and 50 dph males. **d-f.** Representative images of parasagittal slices through the arcopallium of 20, 35, and 50 dph females. In all panels, RA borders are indicated by white arrows. Note the increasing sex difference in RA size across ages, as well as the more pronounced myelination (dark fibers) in RA and surrounding arcopallium in males compared to females. Dorsal is up and anterior to the left.

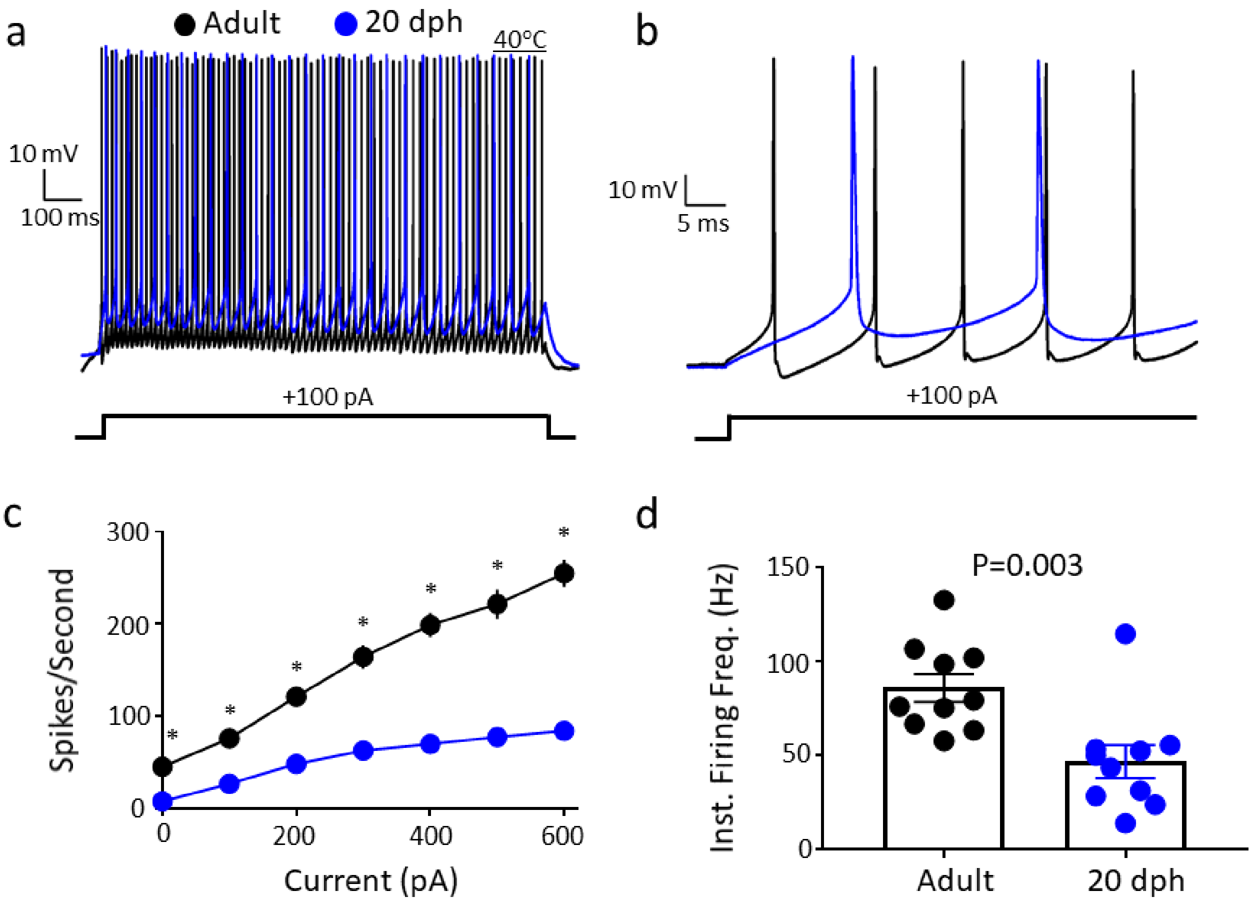


Supplementary Figure 6. Intrinsic excitable properties recorded in whole cell current clamp at 40°C in RAPNs of 20 dph and adult male zebra finches. **a.** Overlay of APs (top) elicited by a +100 pA current injection (bottom) in 20 dph (blue) and adult (black) finches. **b.** Expanded x-axis from (a) showing the first few APs. **c.** Average number of spikes/sec as a function of current injected in 20 dph and adult males (two-way ANOVA with Tukey's post hoc; $P < 0.0001$, $F (6, 110) = 17.62$, N (cells/age) = 8/adult and 10/20 dph finches). Stars depict significant age differences in RA determined by post hoc analyses; * = $P \leq 0.05$. **d.** Average instantaneous firing frequency measured in response to a +100 pA current injection in 20 dph and adult males; N (cells/age) = 10/20 dph and 10/adults; Student's t-test.

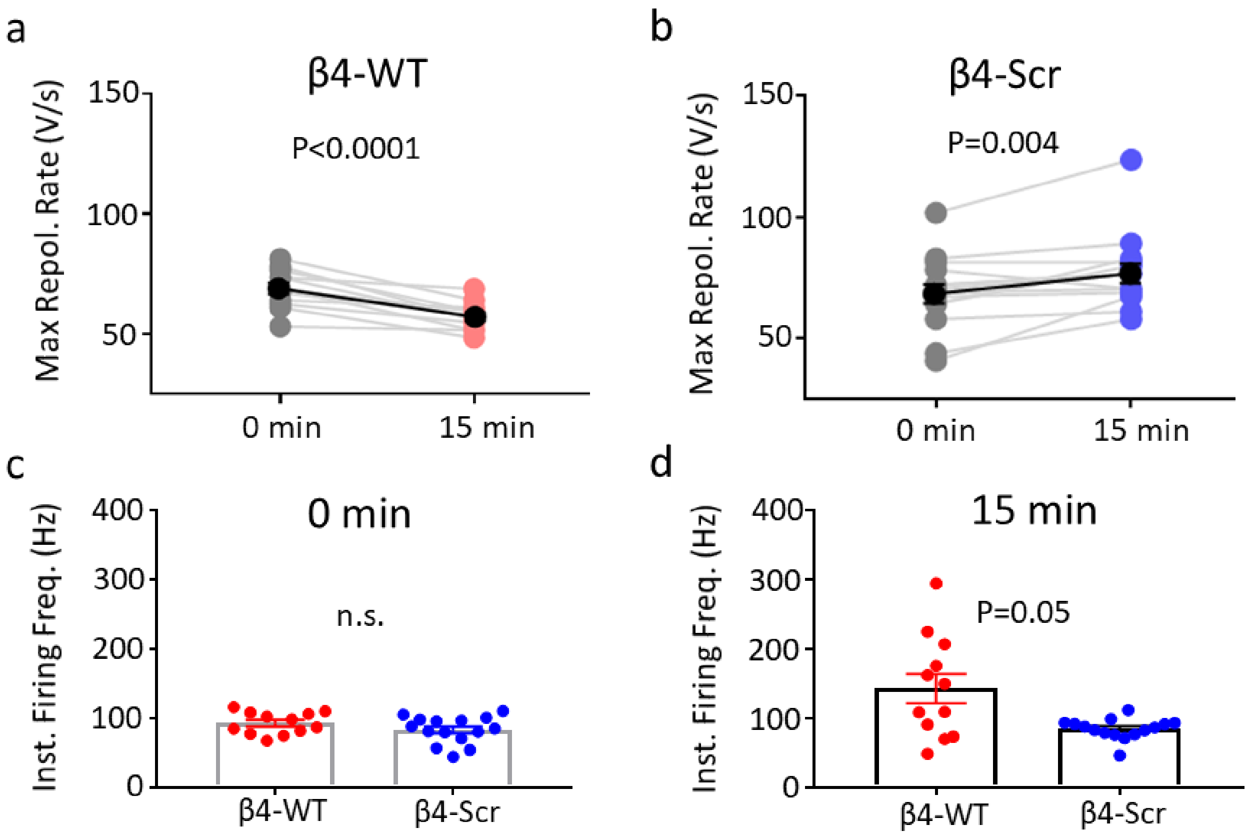


Supplementary Figure 7. Effects of Navβ4 C-terminal peptide (β4-WT) on intrinsic excitable properties of RAPNs in 20 dph males. **a-b.** Time-dependent effects of β4-WT or β4-Scr on the maximum repolarization rate in individual cells (mean ± SEM in black; paired t-test). **c-d.** Comparisons of instantaneous firing frequencies of the first 2 APs elicited immediately or 15 min after achieving the whole-cell configuration between cells exposed to β4-WT (red) or β4-Scr (blue); Student's t-test in (c) and Mann-Whitney test in (d). N= 12 and 15 cells recorded with β4-WT and β4-Scr respectively.

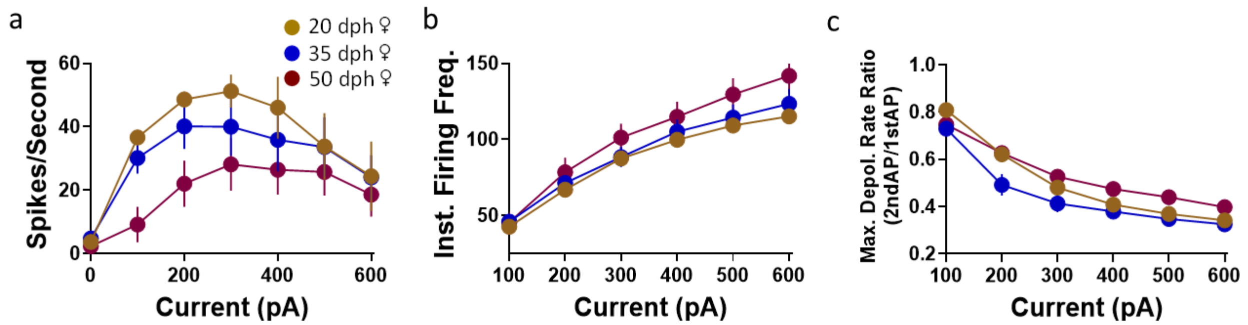


Supplementary Figure 8. Intrinsic excitable properties of RAPNs in female zebra finches across ages. **a.** Average number of spikes/sec as a function of current injected for each age group. Two-way ANOVA with Tukey's post hoc; P = 0.77 , F (12, 296) = 0.6816, N (cells/age) = 21/20 dph, 23/35 dph, 7/50 dph and 18/adult cells. **b.** Average instantaneous firing frequency (IFF) measured as a function of current injected for each age group. Two-way ANOVA with Tukey's post hoc; P = 0.99 , F (10, 211) = 0.3687, N (cells/age) = 21/20 dph, 23/35 dph, 7/50 dph and 18/adult cells. **c.** Fold-change of the maximum depolarization rate from the first to the second AP as a function of current injected for each age group. Two-way ANOVA with Tukey's post hoc; P = 0.51 , F (10, 207) = 0.9226, N (cells/age) = 21/20 dph, 23/35 dph, 7/50 dph and 18/adult cells.

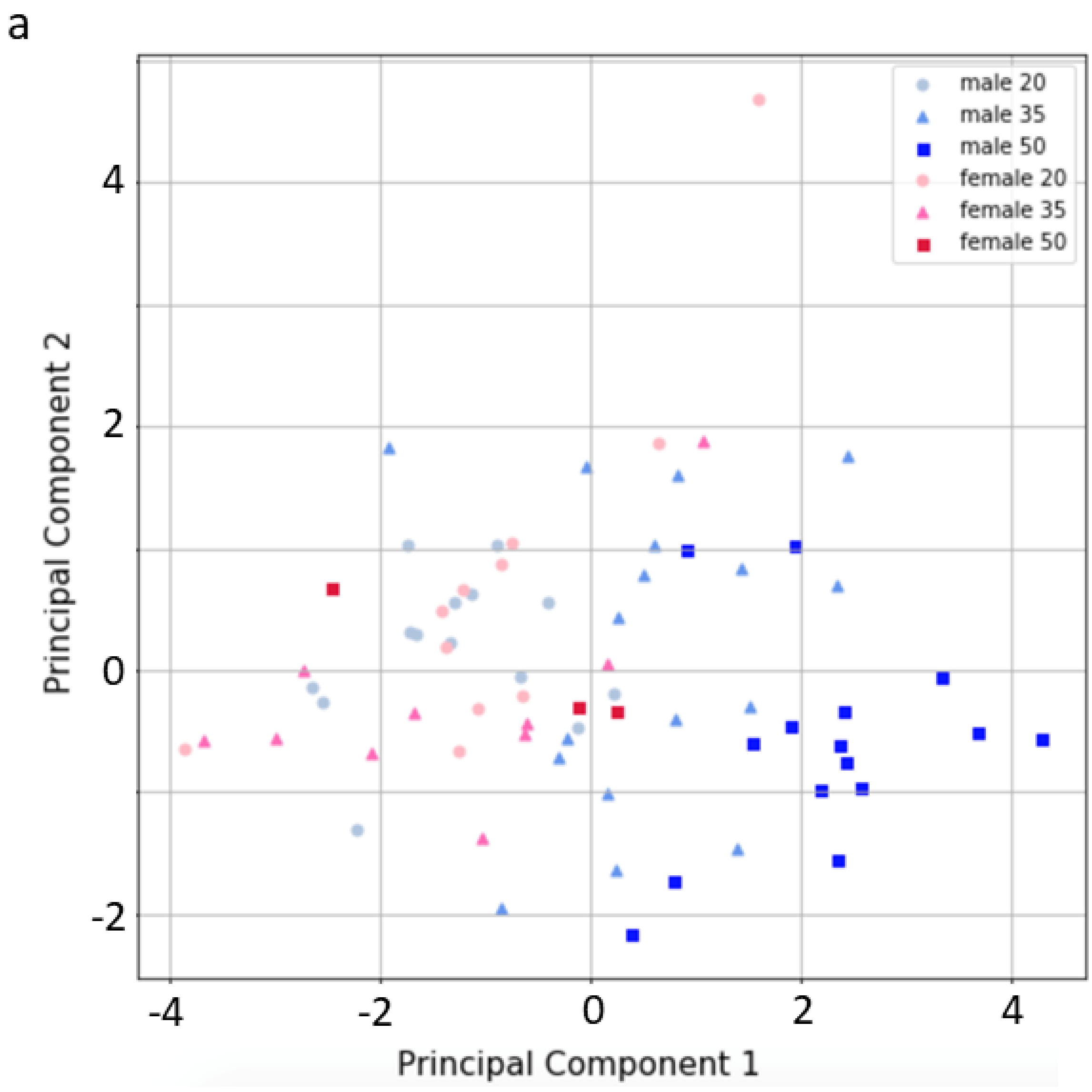


Supplementary Figure 9. Principle component analysis (PCA) for spontaneous APs in female and male finches across development. PCA of spontaneous spike frequency, threshold, half-width, maximum depolarization rate and maximum repolarization rate in males (blue) and females (red), different color ages represented by different shapes and color shades. PC1 and PC2 accounted for 53% and 20%, respectively, of the variance across ages and sex.

### Tables and Legends:

| | Male ♂ | | | | | |
|---|---|---|---|---|---|---|
| **Age (dph)** | **20 (N=18)** | **35 (N=18)** | **50 (N=18)** | **Adult RA (24$^{o}$C; N=18)** | **Adult RA (40$^{0}$C; N=8)** | **Adult (Out) (N=12)** |
| **Input Res. (MΩ)** | 422 ± 32.7$^{\Psi\nabla\Lambda}$ | 285.4 ± 19.9$^{\Phi\nabla\Lambda}$ | 175.7 ± 17.6$^{\Phi\Psi}$ | 190.2 ± 25.4$^{\Phi\Psi}$ | - | 276 ± 36.8$^{\Lambda}$ |
| **Tau (ms)** | 39.0 ± 2.8$^{\Psi\nabla\Lambda}$ | 28. ± 3.5$^{\Phi}$ | 25.9 ± 3.0$^{\Phi}$ | 18.8 ± 1.7$^{\Phi}$ | - | 47.5 ± 4.0$^{\Lambda}$ |
| **Capacitance (pF)** | 103.3 ± 15.8 | 109.0 ± 17.4 | 155.6 ± 19.5 | 114.3 ± 15.3 | - | 195.6 ± 23.3$^{\Lambda}$ |
| **Spont. Freq. (Hz)** | 5.7 ± 0.7 | 5.0 ± 1.4 | 6.8 ± 1.0 | 9.2 ± 2.5 | 45 ± 10.3 | 2.3 ± 1.2$^{\Lambda}$ |
| **% Spont.** | 77.8 | 94.4 | 80.0 | 94.4 | 100 | 41.7 |
| **Thresh. (mV)** | -43.7 ± 0.8$^{\Lambda}$ | -48.1 ± 1.5$^{\Lambda}$ | -48.3 ± 0.7$^{\Lambda}$ | -54 ± 2.0$^{\Phi\Psi\nabla}$ | -44.1 ± 0.7 | -38.3 ± 0.4$^{\Lambda}$ |
| **HW (ms)** | 1.8 ± 7E10-2$^{\Psi\nabla\Lambda}$ | 1.2 ± 8E10-2$^{\Phi\nabla\Lambda}$ | 0.8 ± 5E10-2$^{\Phi\Psi}$ | 0.6 ± 21E-2$^{\Phi\Psi}$ | 0.18 ± 0.02 | 2.1 ± 0.2$^{\Lambda}$ |
| **Amp. (mV)** | 85.1 ± 1.0 | 88.6 ± 1.6 | 90.0 ± 1.8 | 90.5 ± 1.9 | 62.7 ± 3.0 | 62.3 ± 8$^{\Lambda}$ |
| **Max. D. (V/s)** | 260 ± 15.1$^{\Psi\nabla\Lambda}$ | 364.8 ± 20.6$^{\Phi\nabla\Lambda}$ | 488.8 ± 22.1$^{\Phi\Psi}$ | 534.5 ± 33.6$^{\Phi\Psi}$ | 592.4 ± 66.5 | 144 ± 30.8$^{\Lambda}$ |
| **Max. R. (V/s)** | 42.8 ± 2.7$^{\Psi\nabla\Lambda}$ | 77.9 ± 4.8$^{\Phi\nabla\Lambda}$ | 130.4 ± 7.6$^{\Phi\Psi\Lambda}$ | 181.9 ± 10.2$^{\Phi\Psi\nabla}$ | 478.9 ± 63.5 | 41.1 ± 8.2$^{\Lambda}$ |
| **Estim. $I_{Na}$ (nA)** | 26.9 | 39.8 | 76.1 | 61.1 | - | 28.2 |

Table 1. Passive and spontaneously active properties of male arcopallial neurons. A one-way ANOVA with a Tukey post hoc test was used for each measurement inside RA across the four ages indicated. Input Res.- Input resistance, $P < 0.0001$, $F (3, 63)$; Tau- Membrane time constant, $P = 0.0003$, $F (3, 52)$; Capacitance – Membrane capacitance, $P = 0.12$, $F(3, 52)$; Spont. Freq.- Spontaneous firing frequency, $P = 0.26$, $F (3, 68) = 1.379$; % Spont. - proportion of recorded cells that produced spontaneous APs, $P < 0.0001$, $F (3, 63) = 23.15$; Thresh.- AP threshold, $P < 0.0001$, $F (3, 56) = 9.281$; HW- AP half-width, $P < 0.0001$, $F (3, 56) = 57.31$; Amp.- AP amplitude, $P = 0.25$, $F (3, 56) = 1.415$; Max. D.- Maximum depolarization rate, $P < 0.0001$, $F (3, 56) = 22.03$; Max. R.- Maximum repolarization rate, $P < 0.0001$, $F (3, 56) = 62.24$. Estim. $I_{Na}$- Estimated $Na^+$ current during the depolarization phase of spontaneous APs as determined by the *maximum repolarization rate x membrane capacitance*. For comparisons between measurements within RA and outside RA in adults a Student's t-test was used.

$^{\Phi}$ **data is significantly different from 20 dph ($p \leq 0.05$)**

$^{\Psi}$ **data is significant different from 35 dph ($p \leq 0.05$)**

$^{\nabla}$ **data is significant different from 50 dph ($p \leq 0.05$)**

$^{\Lambda}$ **data is significant different from Adult RA ($p \leq 0.05$)**

| | 0.1 mS | | 0.15 mS | | 0.2 mS | | 0.25 mS | |
|---|---|---|---|---|---|---|---|---|
| | **OFF** | **ON** | **OFF** | **ON** | **OFF** | **ON** | **OFF** | **ON** |
| **# APs** | 44.9 ± 3.9 | 43.6 ± 3.9* | 44 ± 4.0 | 39.1 ± 5.1* | 37.4 ± 3.7 | 26.9 ± 4.6 $^{+}$ | 44.2 ± 3.4 | 35 ± 4.37* |
| **Amp. (mV)** | 57.2 ± 3.0 | 44.1 ± 7.3 | 62 ± 2.8 | 52.9 ± 2.8$^{>}$ | 67.5 ± 2.4 | 55.7 ± 2.8$^{+}$ | 65.7 ± 1.8 | 50.4 ± 2.8$^{>}$ |
| **HW (ms)** | 3.2 ± 0.2 | 3.0 ± 0.4 | 2.9 ± 0.7 | 3.1 ± 0.2* | 2.5 ± 0.1 | 2.7 ± 0.2 | 2.6 ± 0.1 | 3.0 ± 0.3 |
| **Max D. (V/s)** | 84.2 ± 9.1 | 67 ± 8.1* | 96.4 ± 9.4 | 72.7 ± 7.0$^{+}$ | 118.2 ± 9.6 | 85.6 ± 8.2$^{>}$ | 109.6 ± 6.6 | 71.9 ± 6.5$^{>}$ |
| **Max R. (V/s)** | 28.7 ± 2.9 | 25.1 ± 2.2* | 30 ± 2.2 | 26.9 ± 2.1$^{+}$ | 32.2 ± 2.3 | 30.1 ± 2.4 | 32 ± 1.4 | 27.6 ± 1.9$^{+}$ |
| **Peak (mV)** | 22.5 ± 3.4 | 18.2 ± 3.2$^{+}$ | 24.9 ± 3.2 | 18.8 ± 3.3$^{<}$ | 28.1 ± 2.6 | 18.7 ± 3.1$^{<}$ | 32.7 ± 2.4 | 25.5 ± 2.9$^{+}$ |
| **AHP (mV)** | -39.7 ± 1.8 | -35.2 ± 3.1* | -40.3 ± 2.5 | -36.8 ± 3.1$^{+}$ | -42.6 ± 1.5 | -37.4 ± 2.0$^{>}$ | -44.2 ± 1.6 | -38.6 ± 2.2$^{+}$ |

Table 2. Average AP properties from juvenile male dynamic clamp experiments. Values at the top of each paired data set (as defined by shading) are conductance values used during dynamic clamping. Values in the OFF column represent values obtained during a +300 pA current injection without dynamic clamping. Values in the ON column represent values obtained during the same current injection with dynamic clamping. For comparisons, a Paired t-test was used for each conductance comparing values obtained before and during dynamic clamping. # APs- the number of APs produced during a 1 sec +300 PA current injection, Amp.- AP Amplitude, HW- AP half-width, Max. D.- Maximum depolarization rate, Max. R.- Maximum repolarization rate, Peak- AP peak, AHP- AP afterhyperpolarization peak. * = $p \leq 0.05$, $^{+}$ = $p \leq 0.01$, $^{>}$ = $p \leq 0.001$, $^{<}$ = $p \leq 0.0001$.

| | 0.25 mS | | 0.5 mS | | 1.0 mS | | 1.5 mS | |
|---|---|---|---|---|---|---|---|---|
| | OFF | ON | OFF | ON | OFF | ON | OFF | ON |
| **# APs** | 69.4 ± 8.7 | 49.2 ± 8.1 | 73.6 ± 7.6 | 72.6 ± 11.4 | 79 ± 8.2 | 76.6 ± 8.9 | 72 ± 10.0 | 67.6 ± 7.9 |
| **Amp. (mV)** | 93.1 ± 2.4 | 99.3 ± 2.9 | 94.6 ± 29.2 | 94.1 ± 2.7 | 91.2 ± 1.7 | 94.3 ± 2.6 | 94.2 ± 1.9 | 95.5 ± 2.2 |
| **HW (ms)** | 0.57 ± 0.02 | 0.58 ± 0.03 | 0.57 ± 0.27 | 0.57 ± 0.03 | 0.5 ± 0.03 | 0.6 ± 0.03 | 0.6 ± 0.03 | 0.6 ± 0.03 |
| **Max D. (V/s)** | 487.54 ± 19.3 | 390.6 ± 32.3 | 519.3 ± 108.5 | 531.3 ± 27.5 | 535.6 ± 24.5 | 548.3 ± 25.7* | 545.8 ± 24.8 | 542.2 ± 38 |
| **Max R. (V/s)** | 191.4 ± 11.4 | 163.4 ± 30.0 | 192.7 ± 54.2 | 195.3 ± 16.4 | 197.1 ± 14.9 | 199.9 ± 15.4 | 199.9 ± 15.9 | 201.8 ± 16.8 |
| **Peak (mV)** | 38.08 ± 1.1 | 38.3 ± 1.4$^{+}$ | 40.5 ± 21.4 | 41.6 ± 2.3 | 40.3 ± 1.2 | 40.6 ± 1.3$^{+}$ | 40.96 ± 1.2 | 20.4 ± 0.9 |
| **AHP (mV)** | -62.7 ± 1.0 | -65.5 ± 1.4 | -63.4 ± 1.0 | -65.1 ± 0.93* | -64 ± 1.0 | -65.8 ± 0.9$^{+}$ | -64.4 ± 0.9 | -67.5 ± 0.7$^{+}$ |

Table 3. Average AP properties from adult male dynamic clamp experiments. Values at the top of each paired data set (as defined by shading) are conductance values used during dynamic clamping. Values in the OFF column represent values obtained during a +300 pA current injection without dynamic clamping. Values in the ON column represent values obtained during the same current injection with dynamic clamping. For comparisons, a Paired t-test was used for each conductance comparing values obtained before and during dynamic clamping. # APs- the number of APs produced during a 1 sec +300 PA current injection, Amp.- AP Amplitude, HW- AP half-width, Max. D.- Maximum depolarization rate, Max. R.- Maximum repolarization rate, Peak- AP peak, AHP- AP afterhyperpolarization peak. * = $p \leq 0.05$, $^{+}$ = $p \leq 0.01$, $^{>}$ = $p \leq 0.001$, $^{<}$ = $p \leq 0.0001$.

| **Female ♀** |
|---|

| Age (dph) | 20 (N=21) | 35(N=18) | 50(N=7) |
|---|---|---|---|
| Input Res. (MΩ) | 442.3 ± 34.2$^{\Psi\nabla}$ | 255.1 ± 26.2$^{\Phi}$ | 262.6 ± 22.5$^{\Phi}$ |
| % Spont. | 52.4 | 43.4 | 42.8 |
| Spont. Freq. (Hz) | 2.5 ± 0.7 | 4.7 ± 1.5 | 5.3 ± 2.1 |
| Thresh. (mV) | -46.5 ± 1.8 | -44.1 ± 1 | -47.9 ± 0.8 |
| HW (ms) | 1.7 ± 7E10-2 | 1.6 ± 9E10-2 | 1.5 ± 2E10-1 |
| Amp. (mV) | 88.2 ± 2.6 | 78.5 ± 2.9 | 78.9 ± 1.9 |
| Max. D. (V/s) | 247.5 ± 12.2 | 210.1 ± 19.6 | 255.8 ± 29.9 |
| Max. R. (V/s) | 49.7 ± 2.5$^{\nabla}$ | 52 ± 4.8$^{\nabla}$ | 63.6 ± 11.5$^{\Phi\Psi}$ |

Table 4. Passive and spontaneously active properties of female arcopallial neurons. A one-way ANOVA with a Tukey post hoc test was used for each measurement across the three ages indicated. Input Res.- Input Resistance, % Spont.- proportion of cells that had spontaneous APs; Spont. Freq.- Spontaneous firing frequency, $P = 0.29$, $F (2, 41) = 1.262$; Thresh.- AP threshold, $P = 0.61$, $F (2, 21) = 0.5043$; HW- AP half-width, $P = 0.54$, $F (2, 21) = 0.6422$, Amp.- AP amplitude, $P= 0.15$, $F (2, 21) = 2.065$; Max. D.- Maximum depolarization rate, $P = 0.42$, $F (2, 21) = 0.9094$; Max. R.- Maximum repolarization rate, $P < 0.0001$, $F (2, 21) = 66.05$.

$^{\Phi}$ **data is significantly different from 20 dph ($p \leq 0.05$)**

$^{\Psi}$ **data is significant different from 35 dph ($p \leq 0.05$)**

$^{\nabla}$ **data is significant different from 50 dph ($p \leq 0.05$)**
